# Transsaccadic feature interactions in multiple reference frames: an fMRIa study

**DOI:** 10.1101/413815

**Authors:** Bianca R. Baltaretu, Benjamin T. Dunkley, Simona Monaco, Ying Chen, J.Douglas Crawford

## Abstract

Transsaccadic integration of visual features can operate in various frames of reference, but the corresponding neural mechanisms have not been differentiated. A recent fMRIa (adaptation) study identified two cortical regions in supramarginal gyrus (SMG) and extrastriate cortex that were sensitive to transsaccadic changes in stimulus orientation (Dunkley et al., 2016). Here, we modified this paradigm to identify the neural correlates for transsaccadic comparison of object orientations in: 1) Spatially Congruent (SC), 2) Retinally Congruent (RC) or 3) Spatially Incongruent (SI)) coordinates. Functional data were recorded from 12 human participants while they observed a grating (oriented 45° or 135°) before a saccade, and then judged whether a post-saccadic grating (in SC, RC, or SI configuration) had the same or different orientation. Our analysis focused on areas that showed a significant repetition suppression (Different > Same) or repetition enhancement (Same > Different) BOLD responses. Several cortical areas were significantly modulated in all three conditions: premotor/motor cortex (likely related to the manual response), and posterior-middle intraparietal sulcus. In the SC condition, uniquely activated areas included left SMG and left lateral occipitotemporal gyrus (LOtG). In the RC condition, unique areas included inferior frontal gyrus and the left lateral BA 7. In the SI condition, uniquely activated areas included the frontal eye field, medial BA 7, and right LOtG. Overall, the SC results were significantly different from both RC and SI. These data suggest that different cortical networks are used to compare pre- and post-saccadic orientation information, depending on the spatial nature of the task.

**Significance Statement:** Every time one makes a saccade, the brain must compare and integrate stored visual information with new information. It has recently been shown that ‘transsaccadic integration’ of visual object orientation involves specific areas within parietal and occipital cortex (Dunkley et al., 2016). Here, we show that this pattern of cortical activation also depends on the spatial nature of the task: when the visual object is fixed relative to space, the eye, or relative to neither space nor the eye, different frontal, parietal, and occipital regions are engaged. More generally, these findings suggest that different aspects of trans-saccadic integration flexibly employ different cortical networks.

## Introduction

Every second humans make several saccades, displacing the retinal image from one fixation point to the next, often requiring the visual system to fuse or compare information between different fixations. The process, whereby visual feature information is retained, spatially updated, and integrated across saccades, is called transsaccadic integration (Irwin, 1996; Melcher & Colby, 2008). Transsaccadic integration has been demonstrated to involve a limited capacity storage mechanism similar to visual working memory (Irwin, 1996; Irwin & Gordon, 1998; Prime, Tsotsos, Keith & Crawford, 2007).

In most natural conditions, transsaccadic integration deals with Spatially Congruent stimuli (Hayhoe et al., 1991; Melcher, 2009; Prime et al., 2006), possibly using mechanisms similar to spatial updating across saccades (Hamker, Zirnsak, Ziesche & Lappe, 2011; Melcher & Colby, 2008; Prime et al., 2011). However, performance can be equal or superior when pre- and post-saccadic stimuli appear at the same retinal location (Golomb, Chun & Mazer, 2008; Golomb & Kanwisher, 2012; Golomb, Pulido, Albrecht, Chun & Mazer, 2010). Moreover, other tasks (such pre/post-saccadic comparisons in a single or unique stimulus) do not require spatial congruence in any frame (Prime et al., 2008; Poth et al. 2015; Poth & Schneider, 2016).

Little is known about the neural mechanisms subserving these different processes (Melcher & Colby, 2008; Cavanagh et al., 2010), but it has been shown that some aspects of transsaccadic integration can be disrupted by transcranial magnetic stimulation (TMS) over posterior parietal cortex (Prime et al., 2008), dorsolateral prefrontal cortex (Tanaka, Dessing, Malik, Prime & Crawford, 2014), and early visual cortex (Malik et al., 2015). Direct evidence for transsaccadic feature integration is scant, but monkey lateral intraparietal cortex (LIP) shows signs of modest spatial updating of rudimentary shape information (Subramanian & Colby, 2014). Demonstrating transsaccadic updating of stimulus features with standard univariate and multivariate fMRI techniques has proven difficult (Lescroart et al., 2016). However, a recent functional magnetic resonance imaging adaptation (fMRIa) paradigm showed transsaccadic interactions between feature orientations in human supramarginal gyrus (SMG; adjacent to the human equivalent to LIP) and extrastriate cortex (Dunkley et al., 2016). These areas were saccade-specific (i.e., they did not show significant feature-specific interactions during fixation), whereas other areas showed the opposite: modulations specific to fixation (Dunkley et al., 2016).

The physiological studies described above employed transsaccadic stimuli that were Spatially Congruent. Other studies have identified retina-fixed mechanisms for visual memory (e.g., Fecteau & Munoz, 2005; Harrison & Tong, 2009; Talsma, White, Mathôt, Munoz & Theeuwes, 2013), and many other studies have looked at feature memory and interactions in the absence of saccades (Averbach & Coriell, 1986; Irwin, 1992; Johnson, Hollingworth & Luck, 2008; Logothetis, Pauls & Poggio, 1995; Motter, 1994; Prime et al., 2007). However, to date, no study has directly compared the mechanisms for transsaccadic feature interactions across situations where the stimuli were spatially arranged to be Spatially Congruent, Retinally Congruent, or neither (Spatially Incongruent).

Here, we investigated the neural mechanisms involved in object orientation in multiple reference frames using an fMRI-adaptation (fMRIa) approach (Grill-Spector & Malach, 2001). Based on previous findings, we hypothesized that integration of object information from 1) objects fixed in space would evoke activation in SMG and extrastriate areas (Dunkley et al., 2016), 2) objects fixed with respect to the retina would be associated with activation in retinally organized regions (i.e., occipital cortex, frontal eye fields), and 3) objects neither fixed in space or with respect to the retina might only recruit object-centered areas (i.e., lateral occipital complex, temporal cortex), unless the brain also compared these separate pre- and post-saccadic images within some ‘dorsal stream’ egocentric spatial scheme (Ungerleider & Haxby, 1994; Crawford et al., 2011; Goodale & Milner, 1992, 2008).

Here, participants were presented with, pre-test stimulus at one of two possible orientations, and then with test stimulus either at the same orientation. Such a protocol is capable of producing both repetition suppression (RS: different > same) or repetition enhancement (RE: same > different) effects in cortical regions sensitive to the manipulated feature (Grill-Spector, Henson & Martin, 2006; Grill-Spector & Malach, 2001; James & Gauthier, 2006; Krekelberg et al., 2006). We tested transsaccadic integration of orientation in three spatial configurations: 1) a Spatially Congruent condition, 2) a Retinally Congruent condition and 3) a Spatially Incongruent condition. Based on a previously published experiment (Dunkley et al., 2016) we expected the Spatially Congruent condition to yield transsaccadic RS in SMG and transsaccadic RE in extrastriate cortex. We then asked: would this activation remain the same independent of spatial condition, or would different cortical areas be recruited for different spatial conditions?

Our results identified several common frontal and parietal regions across all three spatial conditions (perhaps related to the motor response), but other regions showed frame-specific responses. The Spatially Congruent task in the current study yielded a cortical pattern of feature modulations that was generally similar to those reported by Dunkley et al. (2016), but with some differences in lateralization and/or location. In contrast, the Retinally Congruent and Spatially Incongruent conditions produced very different patterns of cortical activation, suggesting that the mechanisms for transsaccadic integration are highly dependent on the spatial nature of the task (i.e., the reference frame in which the object was presented).

## Materials and Methods

### Participants

Participants were graduate students with normal or corrected-to-normal vision. 17 volunteers participated in the study (11 females, 6 males, 21-42 years of age, 16 right-handed) and provided informed consent. They had no history of neurological disorders. The analyses presented in this study are based on 12 of these participants (10 females, 2 males, age 21-42, 11 right-handed individuals). All experiments were approved by our institution’s Human Participants Review Subcommittee.

### Experimental set-up and stimuli

After passing initial MRI safety screening, participants were then informed about the task. Once they understood the task, they lay supine on the MRI table, with their head resting flat within the 32-channel head coil (Figure 1, A). They were then fit with a head-mounted apparatus containing a mirror to reflect images on the screen (within the MRI bore) and an MRI-compatible eye-tracker (i ViewX, SensoMotoric Instruments). This was used to record eye position from the right eye. Behavioral responses were recorded using a ‘button box’ held in the right hand, with the index finger resting on the leftmost button and the middle finger on the adjacent button (Figure 1, A). All button press responses were analysed offline.

**Figure 1.**
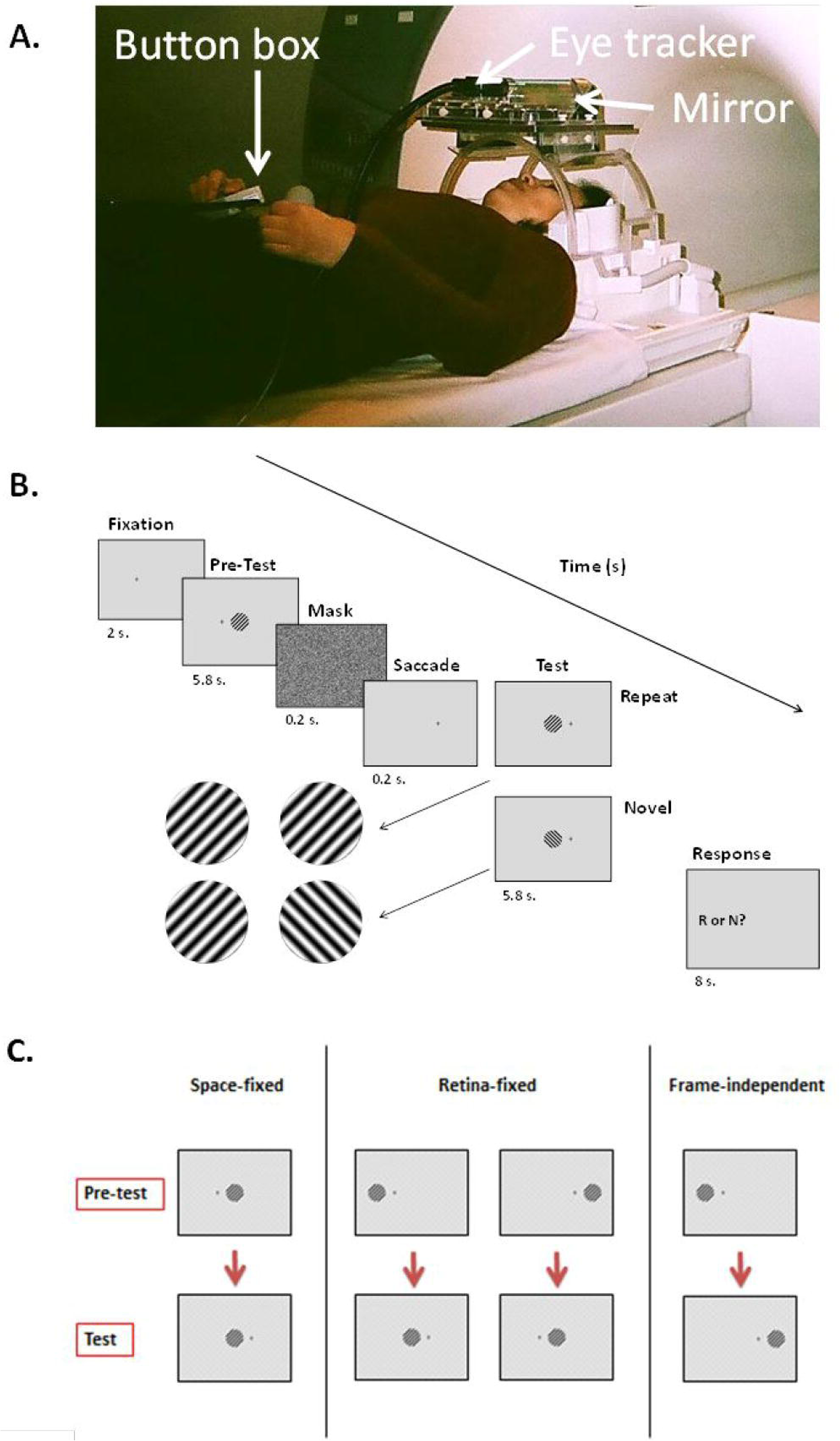
Experimental set-up, paradigm and pre-test, test combinations. ***A***. Experimental set-up showing participant lying supine on MRI table with head mount containing mirror and eye tracker. Button box is held in right hand with index and middle fingers on first two buttons from left to answer during task. ***B***. A typical trial sequence is depicted through time (22 s) for any one of the three spatial condition types (Retinally Congruent, Spatially Congruent or Spatially Incongruent). Stimulus orientations are presented at 45° or 135° and are repeated (‘Repeat’ condition) or are different (‘Novel’ condition). ***C.*** The possible combinations of spatial feature (orientation angle) and location during the first stimulus presentation (‘Pre-test’) and the second stimulus presentation (‘Test’) for the three spatial conditions.

While in the scanner, participants were presented with stimuli consisting of circles 6° in diameter that were filled with a vertical sine-wave grating pattern (0.86 cycles/degree; maximum Michelson contrast) that was averaged to mean luminance of the screen. The grating was presented either at a 45° angle or a 135° angle in standard polar coordinates. The stimuli were presented in the centre of the screen or 12° to the right or left of centre on a light gray background, depending on the spatial condition being tested. The stimuli created with MATLAB (The Mathworks, Inc.) were projected onto a screen in the magnetic bore that was reflected using a mirror onto the participants’ eyes (Figure 1, A).

### Imaging

A 3T Siemens Magnetom TIM Trio magnetic resonance imaging scanner was used to acquire functional data using echo-planar imaging (EPI) sequence (repetition time [TR]= 2000 ms; echo time [TE]= 30 ms; flip angle [FA]= 90°; field of view [FOV]= 192 x 192 mm, matrix size= 64 x 64 with an in-slice resolution of 3 mm x 3 mm; slice thickness= 3mm, no gap) throughout each of the six functional runs in ascending and interleaved order. Thirty-three slices were acquired per volume for a total of 280 volumes of functional data. Within an experimental session, a T1-weighted anatomical reference volume was obtained through an MPRAGE sequence (TR= 1900 ms; FA= 256 mm x 256 mm x 256 mm; voxel size= 1 x 1 x 1 mm^3^). 192 slices were acquired per volume of anatomical data.

### General paradigm/procedure

#### Experiment

We used an fMRIa paradigm to test for adaptation effects to object orientation by presenting a pre-test stimulus at one of two possible orientations (45° or 135°) and a test stimulus whose orientation could be repeated or novel. Each trial (Figure 1, B) started with a 2-s fixation period on a cross at one of two possible positions (6° left or right of centre). This was followed by the presentation of the pre-test stimulus (i.e., the first presentation of the oriented grating) for 5.8 s, after which a 200-ms static random noise mask appeared to prevent retinal afterimages). This was followed by the appearance of a fixation cross at the other fixation cross position for 200 ms prompting participants to make a saccade toward it. The test stimulus was presented for 5.8 s at the same orientation as the pre-test stimulus (‘Repeat’ condition) or at a perpendicular orientation (‘Novel’ condition). The final phase of the trial was a written prompt (‘R or N?’) for 8 s to allow the haemodynamic response to return to baseline as well as to instruct participants to make a response by pressing a button with the right hand to indicate if the orientation of the test stimulus was repeated or novel compared to the pre-test. The first, most-left button was used to indicate that the orientations had been the same (‘Repeat’ condition). The second button was used to indicate that the orientations had been different (‘Novel’ condition).

The presentations of the pre-test and test stimuli were manipulated to depict the spatial conditions probed (Figure 1, C). In the Spatially Congruent condition, the stimuli appeared in the same position on the screen (i.e., in space), but changed relative to the retina upon making the saccade after test presentation (Figure 1, C). This condition was similar to that used in our previous study (Dunkley et al., 2016), except that the diameter of the visual stimulus was reduced to 1/3 the diameter to accommodate the Retinally Congruent and Spatially Incongruent conditions described below, and these three conditions were randomly intermingled during the experiment.

In the Retinally Congruent condition, two possible permutations could be tested: If the fixation cross was initially presented 6° to the left of center, the pre-test stimulus would appear to the left of the fixation cross; then, the fixation cross would subsequently appear 6° to the right of center before the presentation of the test stimulus, which would appear directly in the center of the screen. The opposite could also occur (i.e., the first fixation cross appears 6° to the right of center and the pre-test stimulus appears to the right of the fixation cross; then, the fixation cross moves to 6° to the left of center and the test stimulus appears in the center of the screen). Lastly, in the Spatially Incongruent condition, if the fixation cross appeared 6° to the left of center, the pre-test stimulus would appear to the left of the fixation cross; then, the fixation cross would move to 6° to the right of center and the test stimulus would appear to the right of the fixation cross (and vice versa). As a result, in the Spatially Incongruent condition the pre-test and test stimulus were not matched in space, relative to the retina, or relative to any other reference frame, except that of the object itself. Overall, there were 8 trials for the Spatially Congruent, 8 trials for the Retinally Congruent (to allow for all the permutations that allow the test to end up in the center) and 8 trials for the Spatially Incongruent conditions, for a total of 24 trials in one run, which lasted for ∼ 9 min. in one complete testing sequence, or run. There were an equal number of each trial from each condition, intermingled randomly (through the use of a randomizing MATLAB function) in each run. At the beginning and end of each run, participants were required to fixate centrally for 12 s. There were a total of 6 runs in the entire experiment. This paradigm was modified from Dunkley et al. (2016) and Turi & Burr (2012).

### Analysis

#### Behavioural data

All eye and button press data were analyzed after image acquisition was completed to ensure that the task was being done correctly. Eye position data for each trial were visually inspected to confirm that the subjects fixated on the fixation crosses and performed a saccade at the required times. Any trials in which the eye broke fixation at an inappropriate time were excluded from any further analyses by being designated as confound predictors in fMRI analysis.

Button press responses were also inspected after the experiment. Only trials where button presses correctly indicated the feature condition (Repeat vs. Novel) were included in the analysis. Trials that showed errors in eye and/or button press response were excluded from additional analysis by being designated confound predictors. On this basis of incorrect eye and/or button press responses, the whole data set from one participant who had chance level (50% correct) performance in 5 out of 6 runs was excluded from the analyses. That left 12 participants’ data: three runs were included from one of these participants, who had at least 96 correct trials (88.9% of the total trials), four runs were included from another participant, with at least 125 correct trials (97.7% of the total trials) and five runs were included from a third participant, who had at least 127 correct trials (79.4% of the total trials). The entire data set (all six runs) of the other nine participants were included in analyses, at least 171 correct trials (89.1% of the total trials).

In addition, we employed the same independent localizer used by Dunkley et al. (2016) to identify the parietal eye field. For each trial, participants fixated the centre of a black screen for 2 s, then made self-paced horizontal saccades for 2 s between two fixation crosses placed 12° apart along the middle the screen. Trials were repeated for 256 s (128 volumes) in each subject.

### Functional imaging data: Experimental Task

For each participant, we used a general linear model (GLM) that included 16 predictors. We used a predictor (“Fixate”) for the presentation of the first fixation cross. In the Pre-test phase, we used two predictors: one for a stimulus presented in the left visual field, “Adapt_LVF”, one for a stimulus presented in the right visual field, “Adapt_RVF”. In the test phase, we considered three factors: 3 spatial conditions (Spatially Congruent, Retinally Congruent, Spatially Incongruent) x 2 orientations (Repeat, Novel) x 2 visual hemifields (left: LVF and right: RVF), which resulted in 12 predictors of interest: Spatially Congruent_Repeat_LVF, Spatially Congruent_Repeat_RVF, Spatially Congruent_Novel_LVF, Spatially Congruent_Novel_RVF, Retinally Congruent_Repeat_LVF, Retinally Congruent_Repeat_RVF, Retinally Congruent_Novel_LVF, Retinally Congruent_Novel_RVF, Spatially Incongruent_Repeat_LVF, Spatially Incongruent_Repeat_RVF, Spatially Incongruent_Novel_LVF and Spatially Incongruent_Novel_RVF. We also used a predictor (“Response”) for the button press response. The “Fixate” predictor was 2 s or 1 volume in duration, pre-test predictors were 6 s or 3 volumes in duration, test predictors were 6 s or 3 volumes in duration, and the response predictor was 8 s or 4 volumes long. Predictors were used to generate GLMs for each run for each participant (BrainVoyager QX 2.8, Brain Innovation). A standard haemodynamic response function (HRF; Brain Voyager QX’s default double-gamma HRF) was convolved with the predictor variables, which consisted of a rectangular wave function. The GLMs were then amended to exclude any trials that had incorrect button responses, improper or incorrect eye movement trials and/or trials during which excessive motion was observed. Runs associated with GLMs that had over 50% of the trials excluded were excluded from further functional data analysis.

Functional data for each run for all participants were preprocessed and the motion correction parameters were included as predictors of no interest in the GLM. Preprocessing of functional data involved slice scan time correction (cubic spline), temporal filtering (for removal of frequencies <2 cycles/run) and 3D motion correction (trilinear/sinc). On the basis of motion correction results for each run, data for runs that indicated abrupt movement over 2 mm were excluded from further analyses. The whole data set of four participants was not included in statistical analyses due to head motion exceeding our set threshold. Anatomical data were transformed to a Talairach template (Talairach & Tournoux, 1988). The remaining 12 participants’ functional data were coregistered using gradient-based affine alignment (translation, rotation, scale affine transformation) to the raw anatomical data. Functional data were spatially smoothed using an FWHM of 8 mm.

### Statistical Analyses: Voxelwise

From the final 12 participants’ data, a random effects (RFX) GLM was conducted on the compiled data. The result of the RFX GLM produced a statistical map of activation whereby positive activation showed a repetition suppression (RS) effect (i.e., higher activation in response to ‘Novel’ orientation trials over ‘Repeat’ orientation trials) and negative activation showed a repetition enhancement (RE) effect (i.e., higher activation in response to ‘Repeat’ orientation trials over ‘Novel’ orientation trials). We used a statistical threshold of p<0.05 (t_(11)_=2.25, p=0.04589). To correct for the multiple comparisons problem, a BrainVoyager QX cluster threshold correction plugin was used and applied to each of the three spatial conditions (Forman et al., 1995). Cortical regions that survived cluster threshold correction at p<0.05 (t_(11)_=2.25, p=0.04589) within each of the three spatial conditions were deemed to be statistically significant (cluster threshold for Spatially Congruent condition=131 voxels; cluster threshold for Retinally Congruent condition=172 voxels; and cluster threshold for Spatially Incongruent condition=170 voxels). Contrasts for each condition were applied and β weights were extracted from peak voxels in surviving cortical regions. T-tests were then run on β weights obtained from peak voxels for each of the three conditions to determine if the voxel from the cortical region shows (if at all) an effect specific to the condition or if it is condition-independent. We were able to collapse across visual fields, as there was no statistical difference between visual fields in the conditions. Specifically, pairwise t-tests were performed on Novel vs. Repeat β weights within each area for each condition (i.e., Novel vs. Repeat for Spatially Congruent condition, Novel vs. Repeat for the Retinally Congruent condition, and Novel vs. Repeat for the Spatially Incongruent condition). These t-tests were Bonferroni corrected for multiple comparisons. Only areas that show condition-specific statistically significant effects are described in the results.

The results shown in this manuscript are from the voxelwise analysis of the β weights during the ‘test’ stimulus presentation period, in order to identify areas that show adaptation (RS or RE effects) to orientation in the three spatial conditions tested. From areas that showed such adaptation effects, time courses were then extracted and inspected. Based on inspection of the data, the typical time course was expected to have an initial increase in BOLD activity in response to the first presentation of the stimulus, a second increase in activity in response to the second presentation of the stimulus, and a third increase or plateau in activity during the button press response period. β weight values were also extracted from these areas for further statistical analyses to determine if the observed areas were shared among the spatial conditions or were condition-specific. Areas that are mentioned hereafter met all of the criteria mentioned above. Statistical values (t-statistic and p-values) will be indicated for each of the t-tests run for each brain area (i.e., t_(df)s_ and p_s_ represent the t-statistic and p-value, respectively, for a t-test conducted on the Novel and Repeat β weights within a given area for the Spatially Congruent condition; t_(df)r_ and p_r_ for the Retinally Congruent condition and t_(df)f-i_ and p_f-i_ for the Spatially Incongruent condition).

### Statistical Analyses: Region of Interest

To analyze data from the saccade region localizer, 33 slices were acquired and a total of 128 functional volumes of data were acquired (TR= 2000 ms, TE = 30 ms; in-slice resolution of 3 mm x 3 mm; slice thickness= 3mm, no gap). During anatomical image acquisition and localizer image acquisition, the experimental set-up was kept the same as described above with the exception of the use of an anterior 4-channel head mount and that no button press was required. Functional runs for independent localizers (saccades and retinotopy) were preprocessed in the same way as experimental functional runs are preprocessed (see 2.3b.i). The GLM for the saccade localizer included two predictors: One predictor (“Fixate”) modeled the haemodynamic response for the fixation (2 s), whereas the second predictor (for saccades, “Sacc”) modeled the response for the saccade period (4 s). The fixation-saccade pattern was repeated for a total duration of 256 s. A GLM was created using this protocol file for the saccade localizer. To determine the areas that are involved in self-paced saccades within our tested participants, a fixed effects (FFX) GLM was run on the compiled data of 9 participants’. From this GLM, a contrast was conducted to identify “Sacc” responsive areas.

In order to determine which regions of our data showed Directional selectivity for the left vs. right visual fields, we used a contrast on the experimental runs: Adapt_LVF > Adapt_RVF. The selection criteria based on this contrast was independent from the key experimental contrasts and prevented any bias towards our predictions (Kriegeskorte et al., 2010).

Spatially Congruent localizer: In order to compare the data in the current study with data from Dunkley et al. (2016), we also conducted a region-of-interest (ROI) analysis on the data within each activated activation map from each of the three spatial conditions. To do this, peak voxels (Novel > Repeat) from Dunkley et al. (2016) were obtained for left and right supramarginal gyrus (SMG). Spheres were then created around these peak voxels (257 voxels in left and right SMG) in our data. From the peak voxels of the spheres, β weights were obtained. These were then exported and analysed using SPSS (IBM). The corresponding β -weights were obtained from the Dunkley et al. (2016) study for direct comparison. We also performed a similar procedure using the peak voxel coordinates obtained from the right extrastriate area reported by Dunkley et al. (2016), and the mirror left coordinate (since they did not report any left extrastriate modulation).

Dunkley et al. (2016) suggested that the cortical regions involved in transsaccadic integration for space-fixed stimuli do not necessarily overlap with saccade areas or retinotopic areas. Therefore, we did not use localizers to guide our analysis (with the exception of the comparison with Dunkley et al., 2016 described above), but rather as a means to summarize our main voxelwise results in context of other functional anatomy.

## Results

Overall, the subjects included in this analysis correctly performed the Spatially Congruent Task at 91.4±4.2%, the Retinally Congruent Task at 92.4±3.7%, and the Spatially Incongruent Task at 91.7±4.6% during experimental sessions (mean ± SD across subject, before removal of mistakes for fMRI analysis). There was no significant difference in performance between these tasks (p=0.962). Our objective was to identify and compare areas that show adaptation to object orientation in three spatial conditions. In the following sections, we will first describe the cortical areas that showed repetition suppression (RS) or enhancement (RE) effects derived from each condition separately, compare the Novel vs. Repeat activations of these areas across the different conditions, and finally describe the areas that were common to all spatial conditions. (Please see Table 1 below for a list of the acronyms used in the following results sections, as well as Table 2 for the Talairach coordinates of each cortical area mentioned in the text.)

**Table 1.** List of acronyms of cortical regions mentioned throughout this article and their corresponding, full names.

| Acronym | Full name |
| --- | --- |
| SMG | Supramarginal Gyrus |
| LOtG | Lateral Occipitotemporal Gyrus |
| IBA 7 | Lateral Brodmann Area 7 |
| mBA 7 | Medial Brodmann Area 7 |
| LIP | Lateral Intraparietal Cortex |
| IPS | Intraparietal Sulcus |
| aIPS | Anterior Intraparietal Sulcus |
| mIPS | Middle Intraparietal Sulcus |
| pmIPS | Posterior Middle Intraparietal Sulcus |
| PMd | Doral Premotor Cortex |
| M1 | Primary Motor Cortex |
| Pre-SMA | Presupplementary Motor Area |
| SMA | Supplementary Motor Area |
| PreCS | Precentral Sulcus |
| IFG | Inferior Frontal Gyrus |
| FEF | Frontal Eye Field |

**Table 2.**
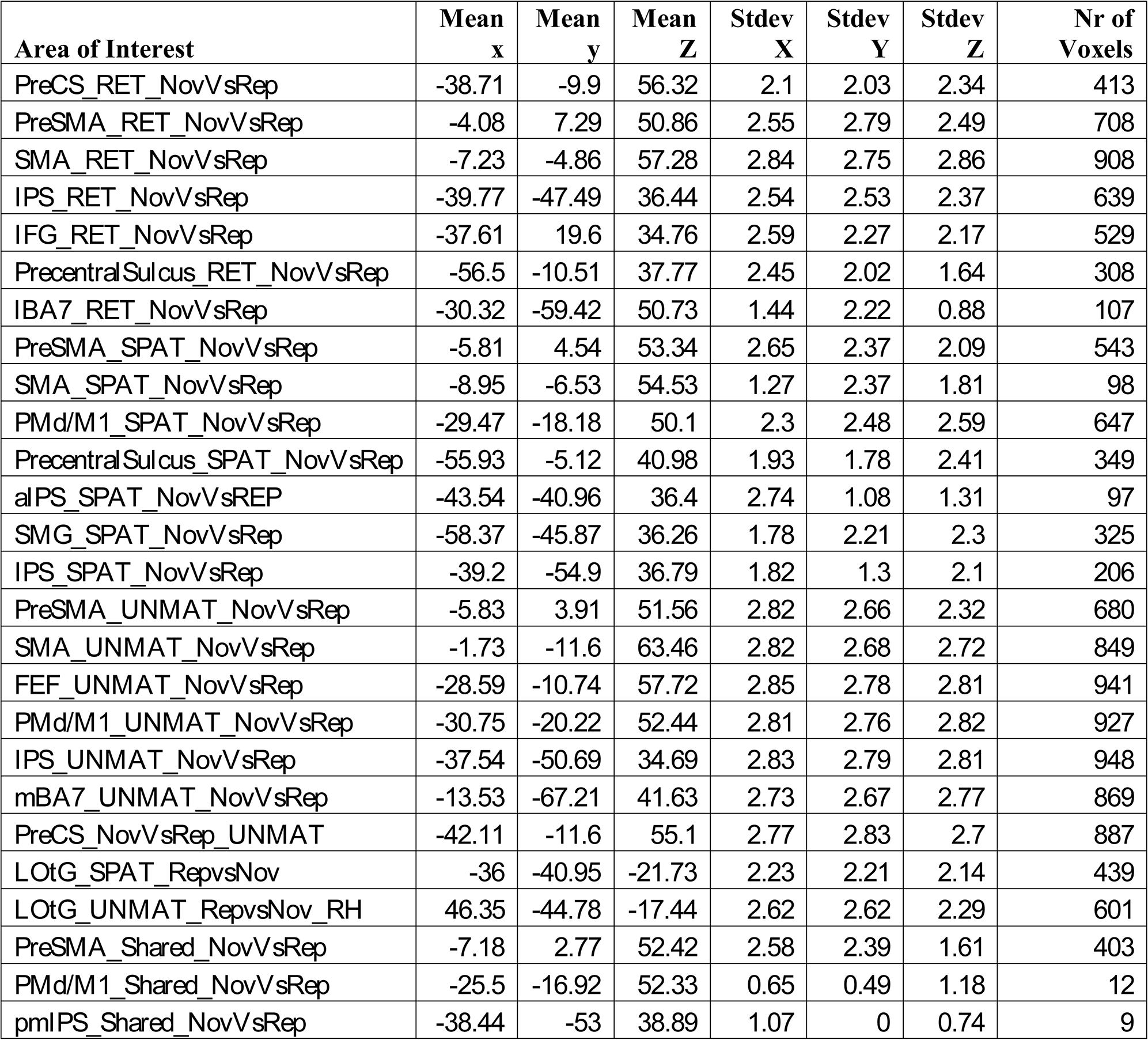
Talairach coordinates of areas of interest resulting from RS or RE effects. In the first column, the cortical region is referred to by its acronym (see Table 1), followed by the spatial condition in which it was found (i.e., RET refers to the Retinally Congruent condition, SPAT refers to the Spatially Congruent condition, and UNMAT refers to the Spatially Incongruent condition), which is subsequently followed by the contrast that was conducted (i.e., NovVsRep refers to the activity in response to Novel (Nov) trials versus Repeat (Rep) trials). If not indicated after the contrast (i.e., NovVsRep), the area was found in the left hemisphere (otherwise, RH refers to an area found within the right hemisphere).

| **Area of Interest** | **Mean**<br>**x** | **Mean**<br>**y** | **Mean**<br>**Z** | **Stdev**<br>**X** | **Stdev**<br>**Y** | **Stdev**<br>**Z** | **Nr of**<br>**Voxels** |
| --- | --- | --- | --- | --- | --- | --- | --- |
| PreCS_RET_NovVsRep | -38.71 | -9.9 | 56.32 | 2.1 | 2.03 | 2.34 | 413 |
| PreSMA_RET_NovVsRep | -4.08 | 7.29 | 50.86 | 2.55 | 2.79 | 2.49 | 708 |
| SMA_RET_NovVsRep | -7.23 | -4.86 | 57.28 | 2.84 | 2.75 | 2.86 | 908 |
| IPS_RET_NovVsRep | -39.77 | -47.49 | 36.44 | 2.54 | 2.53 | 2.37 | 639 |
| IFG_RET_NovVsRep | -37.61 | 19.6 | 34.76 | 2.59 | 2.27 | 2.17 | 529 |
| PrecentralSulcus_RET_NovVsRep | -56.5 | -10.51 | 37.77 | 2.45 | 2.02 | 1.64 | 308 |
| IBA7_RET_NovVsRep | -30.32 | -59.42 | 50.73 | 1.44 | 2.22 | 0.88 | 107 |
| PreSMA_SPAT_NovVsRep | -5.81 | 4.54 | 53.34 | 2.65 | 2.37 | 2.09 | 543 |
| SMA_SPAT_NovVsRep | -8.95 | -6.53 | 54.53 | 1.27 | 2.37 | 1.81 | 98 |
| PMd/M1_SPAT_NovVsRep | -29.47 | -18.18 | 50.1 | 2.3 | 2.48 | 2.59 | 647 |
| PrecentralSulcus_SPAT_NovVsRep | -55.93 | -5.12 | 40.98 | 1.93 | 1.78 | 2.41 | 349 |
| aIPS_SPAT_NovVsRep | -43.54 | -40.96 | 36.4 | 2.74 | 1.08 | 1.31 | 97 |
| SMG_SPAT_NovVsRep | -58.37 | -45.87 | 36.26 | 1.78 | 2.21 | 2.3 | 325 |
| IPS_SPAT_NovVsRep | -39.2 | -54.9 | 36.79 | 1.82 | 1.3 | 2.1 | 206 |
| PreSMA_UNMAT_NovVsRep | -5.83 | 3.91 | 51.56 | 2.82 | 2.66 | 2.32 | 680 |
| SMA_UNMAT_NovVsRep | -1.73 | -11.6 | 63.46 | 2.82 | 2.68 | 2.72 | 849 |
| FEF_UNMAT_NovVsRep | -28.59 | -10.74 | 57.72 | 2.85 | 2.78 | 2.81 | 941 |
| PMd/M1_UNMAT_NovVsRep | -30.75 | -20.22 | 52.44 | 2.81 | 2.76 | 2.82 | 927 |
| IPS_UNMAT_NovVsRep | -37.54 | -50.69 | 34.69 | 2.83 | 2.79 | 2.81 | 948 |
| mBA7_UNMAT_NovVsRep | -13.53 | -67.21 | 41.63 | 2.73 | 2.67 | 2.77 | 869 |
| PreCS_NovVsRep_UNMAT | -42.11 | -11.6 | 55.1 | 2.77 | 2.83 | 2.7 | 887 |
| LOtG_SPAT_RepvsNov | -36 | -40.95 | -21.73 | 2.23 | 2.21 | 2.14 | 439 |
| LOtG_UNMAT_RepvsNov_RH | 46.35 | -44.78 | -17.44 | 2.62 | 2.62 | 2.29 | 601 |
| PreSMA_Shared_NovVsRep | -7.18 | 2.77 | 52.42 | 2.58 | 2.39 | 1.61 | 403 |
| PMd/M1_Shared_NovVsRep | -25.5 | -16.92 | 52.33 | 0.65 | 0.49 | 1.18 | 12 |
| pmlPS_Shared_NovVsRep | -38.44 | -53 | 38.89 | 1.07 | 0 | 0.74 | 9 |

To organize and compare this complex dataset, the following three sub-sections (*Spatially Congruent, Retinally Congruent, and Spatially Incongruent*), Figures 2-5A illustrate cortical regions obtained from the group data for each of these conditions in turn, rendered over an ‘inflated brain’ or two-dimensional anatomic slices from an example participant. These are accompanied by β weights (Figures 2-5B) derived from the peak voxel for the illustrated cortical regions, compared to β weights from the other two conditions at the same coordinates. These sections are then followed by more integrative and comparative descriptions of the data.

**Figure 2.**
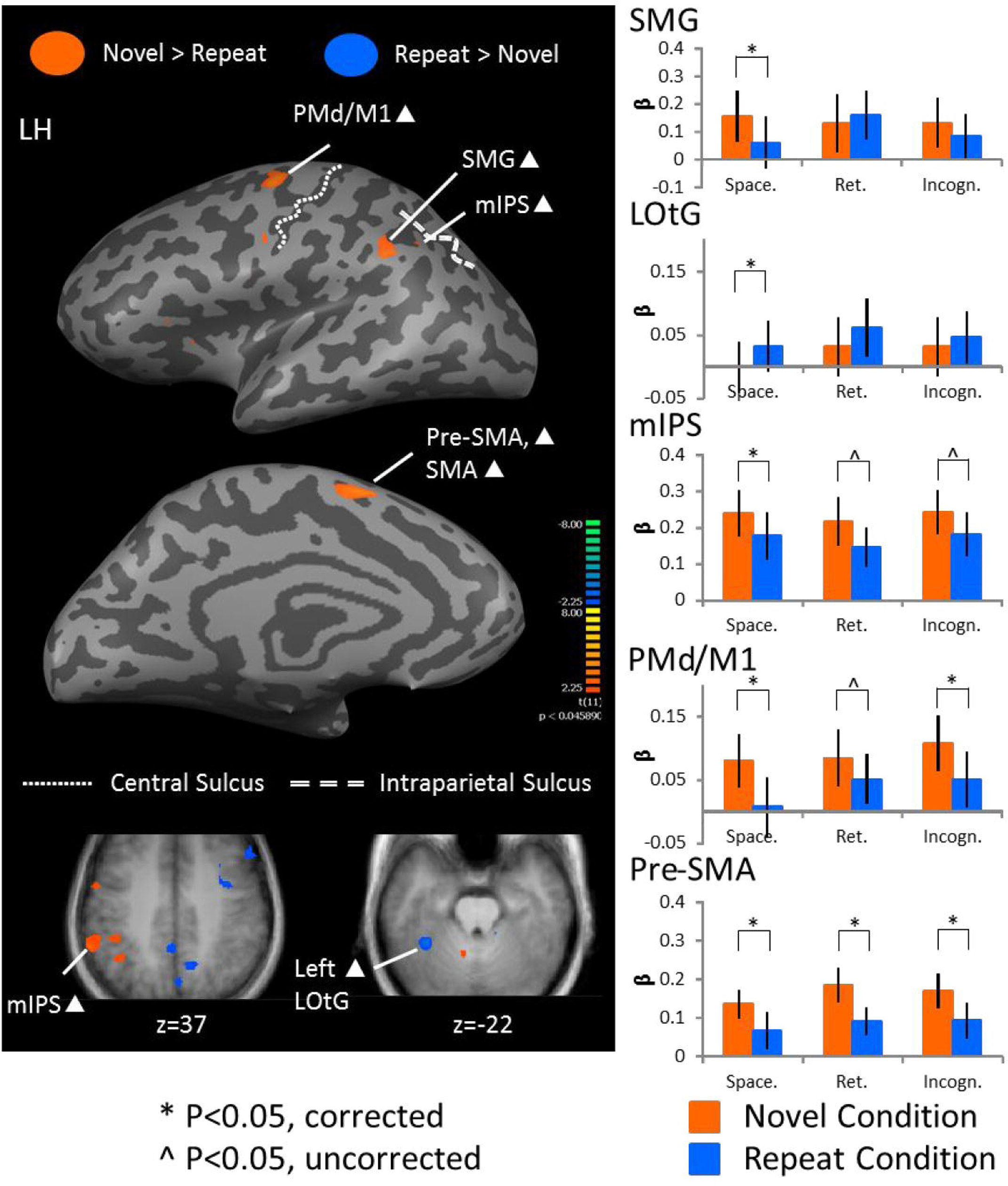
Voxelwise statistical map obtained using an RFX GLM (n=12) for Novel > Repeat in the Spatially Congruent condition (p<0.05). **A:** Activation map is displayed on inflated brain of a representative subject. Activation in orange depicts Repetition Suppression (RS) effects (i.e., where activation is higher for Novel > Repeat), whereas activation in blue depicts Repetition Enhancement (RE) effects (i.e., where activation is higher for Repeat > Novel). Additional slices at bottom show full extent of mIPS and LOtG activation. Areas of interest included SMG: Supramarginal Gyrus, LOtG: Lateral Occipitotemporal Gyrus, mIPS: middle Intraparietal Sulcus, PMd/M1: dorsal Premotor, Primary Motor Cortex, Pre-SMA: Pre-Supplementary Motor Area and SMA: Supplementary Motor Area. **B**: β weights on the right compare peak voxels (which are Bonferroni corrected) from the Spatially Congruent condition to the Retinally Congruent, and Spatially Incongruent conditions within each cortical region. Specificity for the Spatially Congruent condition was found in SMG and LOtG. White triangles indicate cortical regions that passed a p<0.05, but did not pass cluster threshold correction in the left panel. * indicate statistically significant (corrected) differences between Novel and Repeat β weights on the graphs in the right panel, whereas ^ indicate statistically significant differences (uncorrected).

### Spatially Congruent condition

We used a Novel > Repeat contrast for all Spatially Congruent trials: (Spatially Congruent_Novel_LVF + Spatially Congruent_Novel_RVF) > (Spatially Congruent_Repeat_LVF + Spatially Congruent_Repeat_RVF)) to identify brain regions that show adaptation to orientation in a space-fixed condition. The *left panel* of Figure 2 shows an activation map overlaid on an ‘inflated brain’ rendering of an exemplary participant in the top two images and horizontal slices of an average brain of the 12 participants in the bottom two images. Figure 2 (*right column*) shows the corresponding β weights derived from these ‘Spatially Congruent’ areas, and also compares these to the β weights derived from the same areas in the other tasks.

As seen in the left panel of Figure 2, in the Spatially Congruent condition, RS effects (orange) were observed in parietal areas including the supramarginal gyrus (SMG; t_(11)s_= 2.66, p_s_= 0.011, t_(11)r_= -0.58, p_r_= 0.288, t_(11)f-i_= 1.53, p_f-i_=0.077) and the intraparietal sulcus (IPS; t_(11)s_= 3.27, p_s_= 0.004, t_(11)r_= 2.04, p_r_= 0.033, t_(11)f-i_= 2.34, p_f-i_=0.020) in the left hemisphere (SMG in the righthemisphere did not approach significance). We also observed RS effects in left hemisphere frontal motor areas including primary motor cortex (PMd/M1; t_(11)s_= 3.56, p_s_= 0.002, t_(11)r_= 1.82, p_r_= 0.048, t_(11)f-i_= 3.78, p_f-i_=0.002), pre-supplementary motor area (pre-SMA, SMA; t_(11)s_=3.43, p_s_= 0.003, t_(11)r_= 2.75, p_r_= 0.009, t_(11)f-i_= 2.80, p_f-i_= 0.009) and SMA(t_(11)s_= -2.68, p_s_= 0.011, t_(11)r_= -3.32, p_r_= 0.003, t_(11)f-i_= -2.70, p_f-i_= 0.010). RE effects (blue) were observed in the lateral occipitotemporal gyrus (LOtG; t_(11)s_= 4.44, p_s_= 0.0005, t_(11)r_= 1.16, p_r_= 0.135, t_(11)f-i_= 1.23, p_fi_=0.122) in the left hemisphere. (For all of the areas mentioned here, the associated Talairach coordinates may be found in Table 2.)

Turning to the β weights for these areas in the right column of Figure 2, there was (by definition) always a significant difference between Novel and Repeat conditions in the spatial condition (from which these regions were derived), but this was not necessarily the case for the other spatial conditions. SMG and LOtG showed Spatially Congruent-specific results, whereas the RS effects in mIPS, PMd/M1 and pre-SMA were common to all three conditions. Overall, our Spatially Congruent condition produced very specific feature-specific modulations in the left hemisphere, with RS effects in parietal and frontal areas, whereas RE effects were in occipitotemporal regions.

The inferior parietal and occipitotemporal results of this experiment were reminiscent of those of Dunkley et al. (2016), including repetition suppression in SMG and repetition enhancement in visual association cortex, with two key differences: our results here were lateralized to the left hemisphere (as opposed to the right hemisphere), and our extrastriate area was localized more anterior-lateral, bordering on temporal cortex (as opposed to an area overlapping human V4). This may have been due to stimulus differences, which include: 1) 6° gratings here compared to 18° gratings in Dunkley et al. (2016); 2) fixation crosses spanned 6° here, whereas they spanned 27° in Dunkley et al. (2016); and 3) stimuli came within 3° of the outer edge of the screen here, in contrast to within 9° in Dunkley et al. (2016). To perform a more direct comparison between these two studies, we located the peak voxel contrast (Novel – Repeat) within two spheres created around the left and right SMG voxel coordinates reported by Dunkley et al. (Figure 3A), and obtained β weight values for these voxels (Figure 3 B). T-tests were then performed on the Novel vs. Repeat condition. Consistent with our voxelwise analysis above, there was a statistically significant difference between Novel and Repeat in the left SMG (p=0.011), with no difference in right SMG (p=0.464). A similar procedure was performed using the peak voxel coordinate from the right extra-striate area reported by Dunkley et al. (2016), and its mirror left location, but this yielded no significant results in our data (not shown).

**Figure 3.**
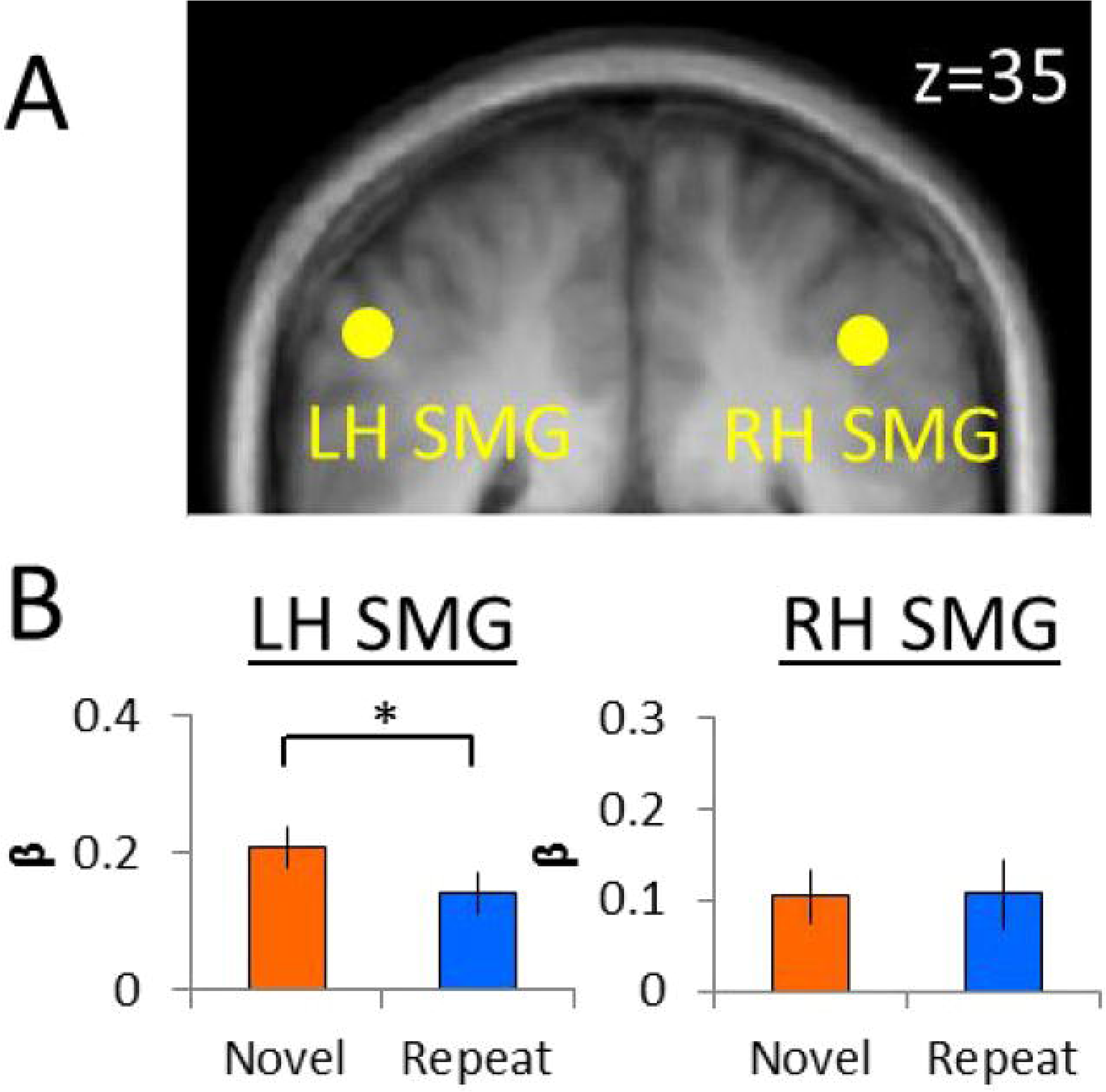
Region-of-interest analysis results. Voxelwise statistical map obtained using an RFX GLM (n=12) for Novel > Repeat in the Spatially Congruent condition (p<0.05). **A**: Left and right SMG are depicted using the yellow circles on a coronal slice through an average brain of all of the participants in the analysis. **B**: β weights obtained from peak voxels within each labeled region and compared (and Bonferroni corrected) between the spatial conditions. β weights are shown for Novel (orange) and Repeat (blue) trials for each of the spatial conditions in the left SMG (*left*) and in the right SMG (*right*). * indicates statistically significant (corrected) differences between Novel and Repeat β weights.

### Retinally Congruent condition

Retinally Congruent trials, a Novel > Repeat contrast was used: Retinally Congruent_Novel_LVF + Retinally Congruent_Novel_RVF) > (Retinally Congruent_Repeat_LVF + Retinally Congruent_Repeat_RVF) to identify areas showing adaptation to orientation in a retinally Congruent condition. Figure 4 shows activation maps overlaid on an inflated brain (*left panel*) and bar graphs of β weights (*right column*) from Retinally Congruent trials. RS effects were observed in parietal areas such as the IPS (IPS; t_(11)s_= 2.20, p_s_= 0.025, t_(11)r_= 2.38, p_r_= 0.018, t_(11)f-i_= 2.81, p_f-i_=0.009) and lateral BA 7 (lBA 7; t_(11)s_= -1.08, p_s_= 0.153, t_(11)r_= -2.65, p_r_= 0.011, t_(11)f-i_= -1.79, p_f-i_= 0.051) in the left hemisphere. Areas in frontal cortex, including M1 (t_(11)s_= 1.59, p_s_= 0.070, t_(11)r_= 2.75, p_r_= 0.009, t_(11)f-i_= 2.65, p_f-i_= 0.011), pre-SMA (t_(11)s_= 1.98, p_s_= 0.036, t_(11)r_= 2.78, p_r_= 0.009, t_(11)f-i_= 2.49, p_f-i_= 0.015), SMA (t_(11)s_= -2.15, p_s_= 0.027, t_(11)r_= -3.63, p_r_= 0.002, t_(11)f-i_= -3.11, p_f-i_= 0.005), precentral sulcus (preCS; t_(11)s_= 0.536, p_s_= 0.301, t_(11)r_= 2.16, p_r_= 0.027, t_(11)f-i_= 2.97, p_f-i_= 0.006) and inferior frontal gyrus (IFG; t_(11)s_= 0.403, p_s_= 0.347, t_(11)r_= 3.10, p_r_= 0.005, t_(11)f-i_= 1.39, p_f-i_= 0.096) in the left hemisphere also showed RS effects. RE effects were not observed in this spatial condition. Our β weight analysis suggests that lBA 7, and IFG, showed RS effects specific to the Retina-fixed task, whereas preCS, aIPS, and again PMd/M1/PreSMA/SMA (not repeated here) showed significant feature modulations across all three tasks. In summary, our Retinally Congruent task only produced RS effects in the left parietal and frontal cortex, but involved a slightly larger, more continuous swath of cortical tissue (compared to Spatially Congruent) running from IPS to IFG.

**Figure 4.**
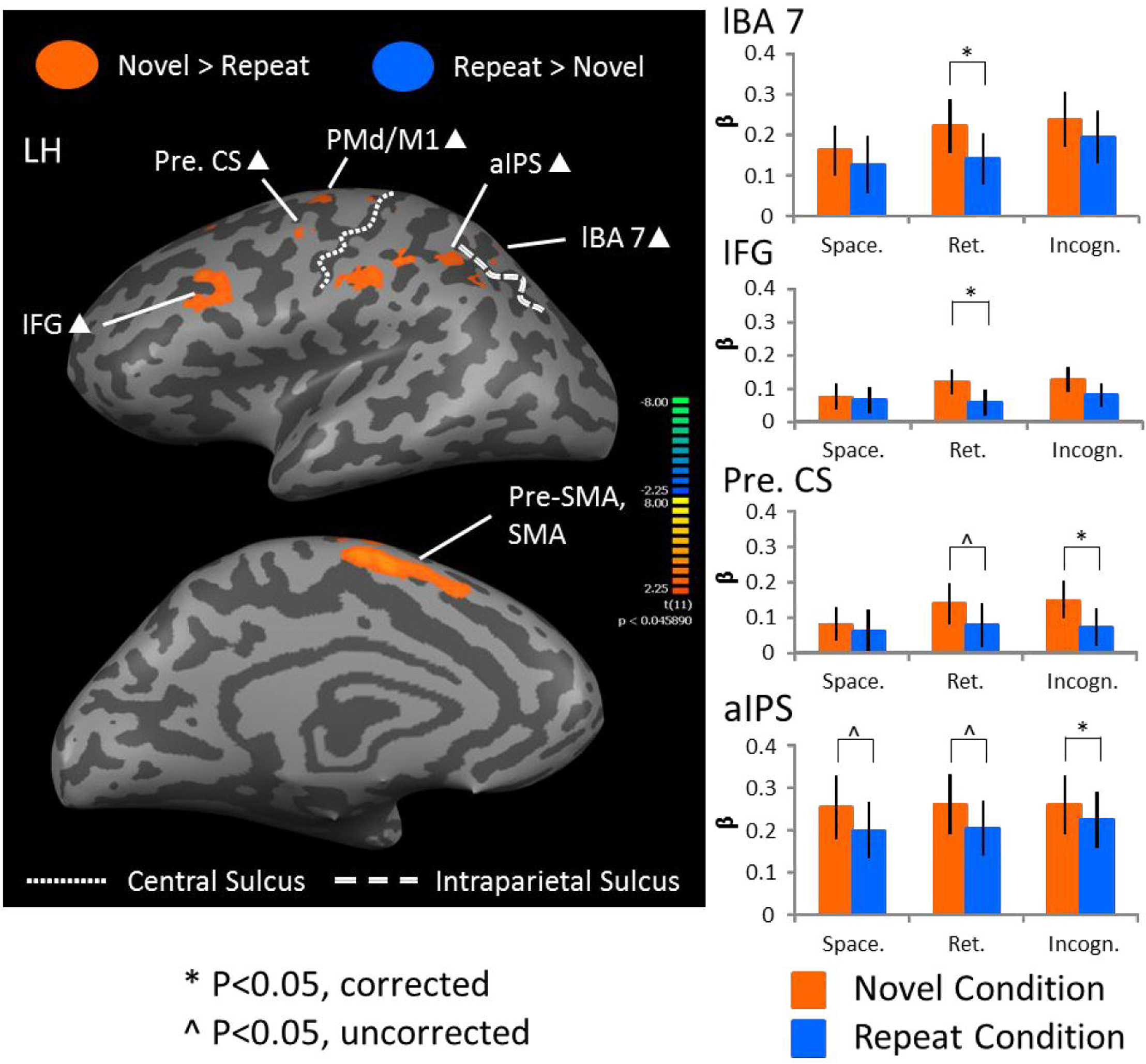
Voxelwise statistical map obtained using an RFX GLM (n=12) for Novel > Repeat in the Retinally Congruent condition (p<0.05). **A**: Activation map is displayed on inflated brain of a representative subject. Activation in orange depicts Repetition Suppression (RS) effects (i.e., where activation is higher for Novel > Repeat), whereas activation in blue depicts Repetition Enhancement (RE) effects (i.e., where activation is higher for Repeat > Novel). Areas of interest included lBA 7: lateral Brodmann area 7, aIPS: anterior Intraparietal Sulcus, PMd/M1, Pre. CS: Precentral Sulcus, IFG: Inferior Frontal Gyrus, Pre-SMA and SMA. **B**: β weights on the right compare peak voxels (which are Bonferroni corrected) from the Retinally Congruent condition to the Spatially Congruent, and Spatially Incongruent conditions for each cortical region. We found specificity for the Retinally Congruent condition in l BA 7 and IFG. White triangles indicate cortical regions that passed a p<0.05, but did not pass cluster threshold correction in the left panel. * indicate statistically significant (corrected) differences between Novel and Repeat β weights on the graphs in the right panel, whereas ^ indicate statistically significant differences (uncorrected).

### Spatially Incongruent condition

In the Spatially Incongruent condition, a Novel > Repeat contrast was used (i.e., (Spatially Incongruent_Novel_LVF + Spatially Incongruent_Novel_RVF) > (Frame-independent _Repeat_LVF + Spatially Incongruent_Repeat_RVF)) to identify brain regions that show adaptation to orientation. Activation maps are overlaid onto an inflated brain/horizontal brain slice and on an average brain and bar graphs of β weights are shown (Figure 5, left and right, respectively). RS effects were observed in parietal areas such as IPS (t_(11)s_= 2.65, p_s_= 0.011, t_(11)r_= 2.13, p_r_= 0.028, t_(11)f-i_= 2.63, p_f-i_= 0.012) and medial BA 7 (mBA 7; t_(11)s_= 0.825, p_s_= 0.213, t_(11)r_= 0.729, p_r_= 0.241, t_(11)f-i_= 2.91, p_f-i_= 0.007) in the left hemisphere. Additionally, RS effects were observed in frontal areas such as preCS (t_(11)s_= 0.607, p_s_= 0.278, t_(11)r_= 1.89, p_r_= 0.042, t_(11)f-i_= 3.55, p_f-i_= 0.002), M1 (t_(11)s_= 3.39, p_s_= 0.003, t_(11)r_= 2.05, p_r_= 0.032, t_(11)f-i_= 4.74, p_f-i_= 0.0003), pre-SMA (t_(11)s_= 3.26, p_s_= 0.004, t_(11)r_= 2.74, p_r_= 0.01, t_(11)f-i_= 2.85, p_f-i_= 0.008), SMA (t_(11)s_= -0.807, p_s_= 0.218, t_(11)r_= -1.59, p_r_= 0.070, t_(11)f-i_= -3.97, p_f-i_= 0.001) and frontal eye fields (FEF; t_(11)s_= 1.16, p_s_= 0.135, t_(11)r_= 1.10, p_r_= 0.148, t_(11)f-i_= 3.91, p_f-i_= 0.001) in the left hemisphere. On the other hand, RE effects were observed in LOtG (t_(11)s_= 0.128, p_s_= 0.450, t_(11)r_= -0.094, p_r_= 0.463, t_(11)f-i_= 4.18, p_f-i_= 0.0008) in the right hemisphere. Here, mBA 7, FEF and LOtG showed Spatially Incongruent condition-specific effects. Overall, RS effects in this condition are found within a parietofrontal network (in this case, more widespread network than either of the preceding tasks), whereas RE effects are depicted in right occipital cortex.

**Figure 5.**
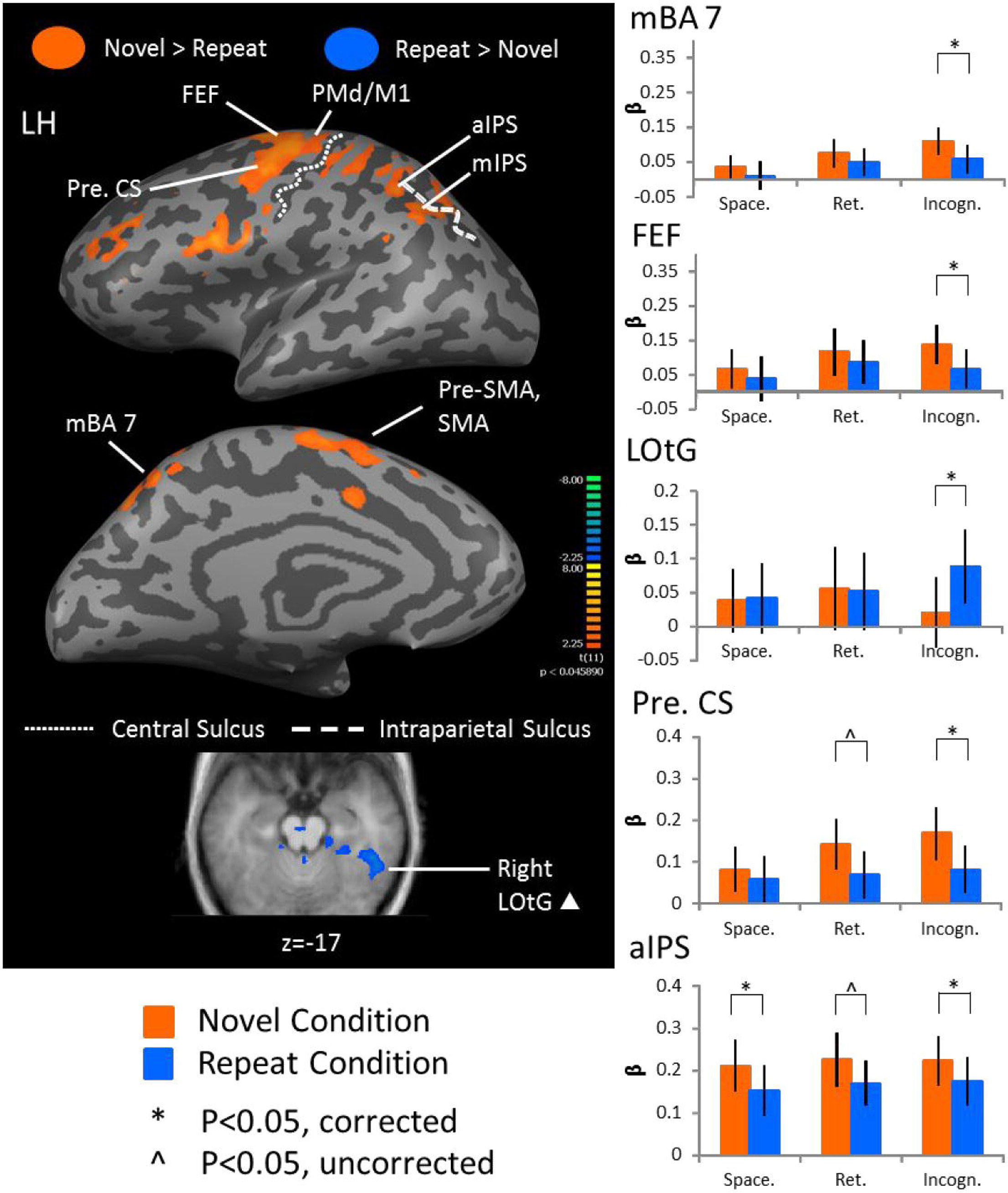
Voxelwise statistical map obtained using an RFX GLM (n=12) for Novel > Repeat in the Spatially Incongruent condition (p<0.05). **A**: Activation map is displayed on inflated brain of a representative subject. Activation in orange depicts Repetition Suppression (RS) effects (i.e., where activation is higher for Novel > Repeat), whereas activation in blue depicts Repetition Enhancement (RE) effects (i.e., where activation is higher for Repeat > Novel). An additional slice at bottom shows full extent of LOtG activation. Areas of interest included SPL, mIPS, aIPS, PMd/M1, FEF: Frontal Eye Field, Pre. CS, Pre-SMA, SMA and LOtG. **B**: β weights on the right compare peak voxels (which are Bonferroni corrected) from the Spatially Incongruent condition to the Spatially Congruent condition, and the Retinally Congruent conditions within each cortical region. We found specificity for the Spatially Incongruent condition in mBA 7 (medial BA 7), FEF and LOtG. White triangles indicate cortical regions that passed a p<0.05, but did not pass cluster threshold correction in the left panel. * indicate statistically significant (corrected) differences between Novel and Repeat β weights on the graphs in the right panel, whereas ^ indicate statistically significant differences (uncorrected).

### ANOVA results

To provide an overall statistical comparison of the three spatial conditions, we a univariate analysis-of-variance (ANOVA) on all of the data from the cortical regions that were obtained from the three contrasts described above. This was done using statistical program R (Core Team, 2013) using an RFX GLM with the following factors: 1) Adaptation (RS or RE), 2) Spatial Condition (Spatially Congruent, Retinally Congruent or Spatially Incongruent), 3) Cortical Region (there were 19 cortical regions in total) and 4) Participant (to account for intersubject variability). ANOVA results indicated that there was a significant main effect of: 1) Adaptation (F_1,1335_=29.529, p=6.54 x 10^-8), 2) Spatial Condition (F_2,1335_=7.148, p=0.000817), 3) Cortical Region (F_18,1335_=12.880, p<2 x 10^-16), and 4) Participant (F_11,1335_=66.909, p=2 x 10^-6). Post-hoc pairwise Tukey t-tests were conducted on the Spatial Condition factor. Results showed that there is a significant difference between the Spatially Congruent condition and the Retinally Congruent condition (p=0.003771) and between the Spatially Congruent condition and the Spatially Incongruent condition (p=0.0025859), but not between the Retinally Congruent and Spatially Incongruent conditions (p=0.9933757). Overall, results indicate that there is a significant difference between: 1) Novel and Repeat condition β weights, 2) Spatially Congruent condition β weights and Retinally Congruent condition and Spatially Incongruent condition β weights, 3) cortical region β weights and 4) participants.

### Conjunction results

Conjunction analyses were used to identify cortical regions that are feature-sensitive and common to all three spatial conditions tested here. Novel > Repeat and Repeat > Novel contrasts were used and subsequently, two condition and three condition conjunctions were conducted to identify the extent of overlap with any of the two-condition conjunctions, as the three-way conjunction results identified small areas of activation showing overlap in all three spatial conditions. Activation maps are shown on an average brain (Figure 6). RS effects were common in motor areas in the left hemisphere such as PMd/M1 (t_(11)s_= 3.02, p_s_= 0.006, t_(11)r_= 2.53, p_r_= 0.014, t_(11)f-i_= 3.08, p_f-i_= 0.005) and pre-SMA (t_(11)s_= -3.59, p_s_= 0.002, t_(11)r_= -2.70, p_r_= 0.010, t_(11)f-i_= -2.65, p_f-i_= 0.011). In the parietal cortex in the left hemisphere, a very small portion of the posterior portion of mIPS (pmIPS; t_(11)s_= 3.04, p_s_= 0.006, t_(11)r_= 2.25, p_r_= 0.023, t_(11)f-i_= 2.23, p_f-i_= 0.024), showed a common RS effect in all three spatial conditions.

**Figure 6.**
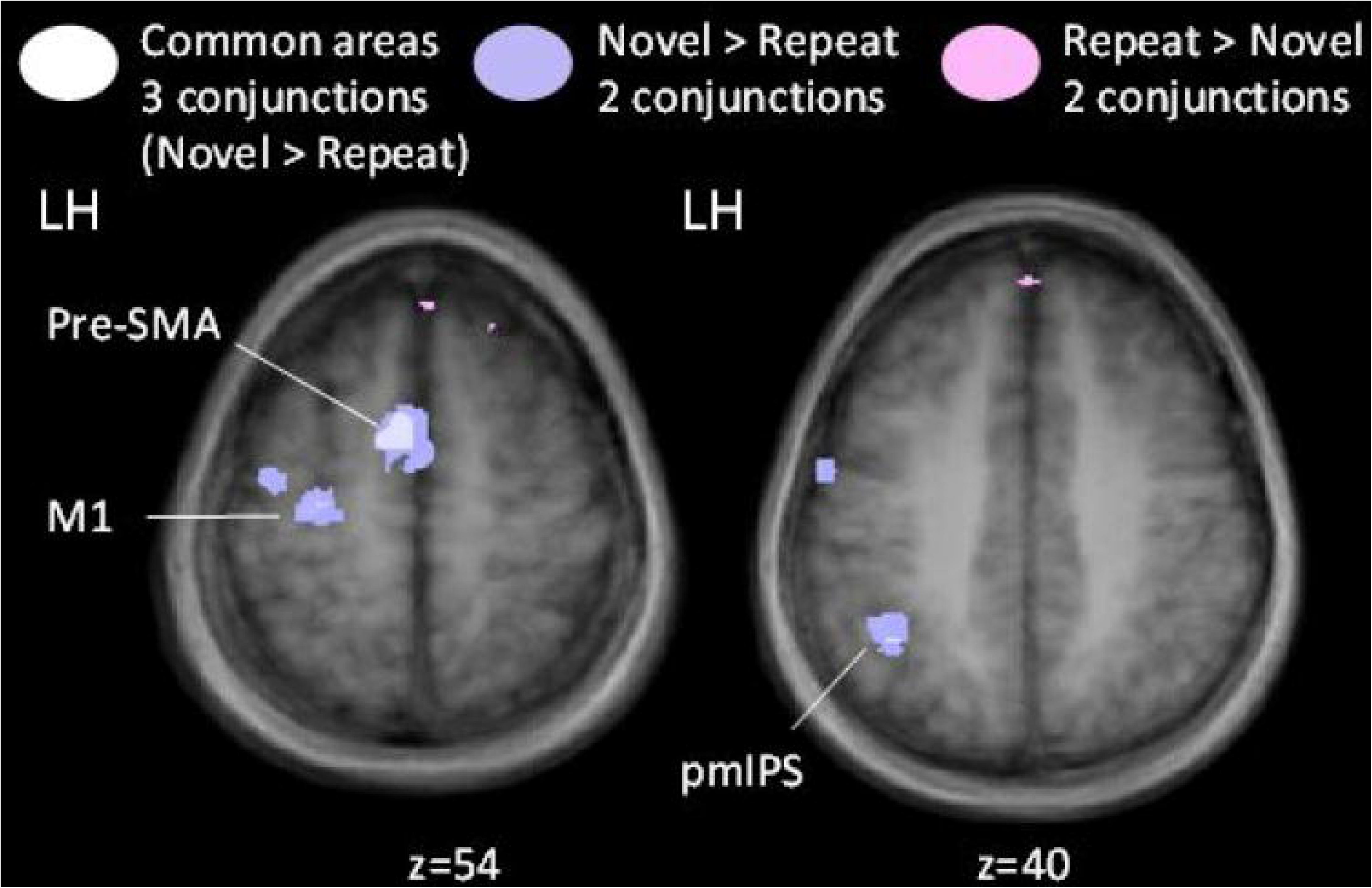
Voxelwise statistical map obtained from an FFX GLM (n=12) for Novel > Repeat in a multiple conjunction analysis. Areas depicted in purple show any two conjunctions using Novel > Repeat contrast (i.e., Spatially Congruent U Retinally Congruent, Spatially Congruent U Spatially Incongruent and/or Retinally Congruent U Spatially Incongruent). Similar conjunctions are shown in pink for a Repeat > Novel contrast. Three-way conjunction analysis results are shown in white for a contrast of Novel > Repeat (i.e., Spatially Congruent U Retinally Congruent U Spatially Incongruent). The specific contrasts used are two-condition and three-condition conjunctions (two-way: 1) (Spatially Congruent_Novel_LVF + Spatially Congruent_Novel_RVF) > (Spatially Congruent_Repeat_LVF + Spatially Congruent_Repeat_RVF) U (Retinally Congruent_Novel_LVF + Retinally Congruent_Novel_RVF) > (Retinally Congruent_Repeat_LVF + Retinally Congruent_Repeat_RVF); 2) (Spatially Congruent_Novel_LVF + Spatially Congruent_Novel_RVF) > (Spatially Congruent_Repeat_LVF + Spatially Congruent_Repeat_RVF) U (Spatially Incongruent_Novel_LVF + Spatially Incongruent_Novel_RVF) > (Spatially Incongruent_Repeat_LVF + Spatially Incongruent_Repeat_RVF); 3) (Retinally Congruent_Novel_LVF + Retinally Congruent_Novel_RVF) > (Retinally Congruent_Repeat_LVF + Retinally Congruent_Repeat_RVF) U (Spatially Incongruent_Novel_LVF + Spatially Incongruent_Novel_RVF) > (Spatially Incongruent_Repeat_LVF + Spatially Incongruent_Repeat_RVF); three-way: (Spatially Congruent_Novel_LVF + Spatially Congruent_Novel_RVF) > (Spatially Congruent_Repeat_LVF + Spatially Congruent_Repeat_RVF) U (Retinally Congruent_Novel_LVF + Retinally Congruent_Novel_RVF) > (Retinally Congruent_Repeat_LVF + Retinally Congruent_Repeat_RVF) U (Spatially Incongruent_Novel_LVF + Spatially Incongruent_Novel_RVF) > (Spatially Incongruent_Repeat_LVF + Spatially Incongruent_Repeat_RVF)).

### Time courses

Figure 7 shows representative time courses derived from selected regions extracted from the previous analyses, all from the left hemisphere (LH) including pmIPS as a common area (A), SMG and LOtG as unique ‘Spatially Congruent’ areas (B), lBA 7 as a unique ‘Retinally Congruent’ area (C), and mBA 7 and FEF as unique ‘Spatially Incongruent’ areas (D). In most cases, one can detect an initial peak BOLD response late in the pre-test phase (6 s after presentation of the test stimulus). This is followed by a second peak BOLD response at the end of the test phase, 4-6 s after presentation of the test stimulus. This is then followed by a third peak (probably related to the behavioral response) about 4 s after the cue to respond. Importantly, the initial separation between Novel (orange) and Repeat (blue) conditions coincides with the second visual peak – as expected if this is related to early processing of the difference – and continues approximately until the peak of the behavioral response. This is why these areas emerge from our analysis of the GLM/β weights locked to the 2^nd^, test stimulus.

**Figure 7.**
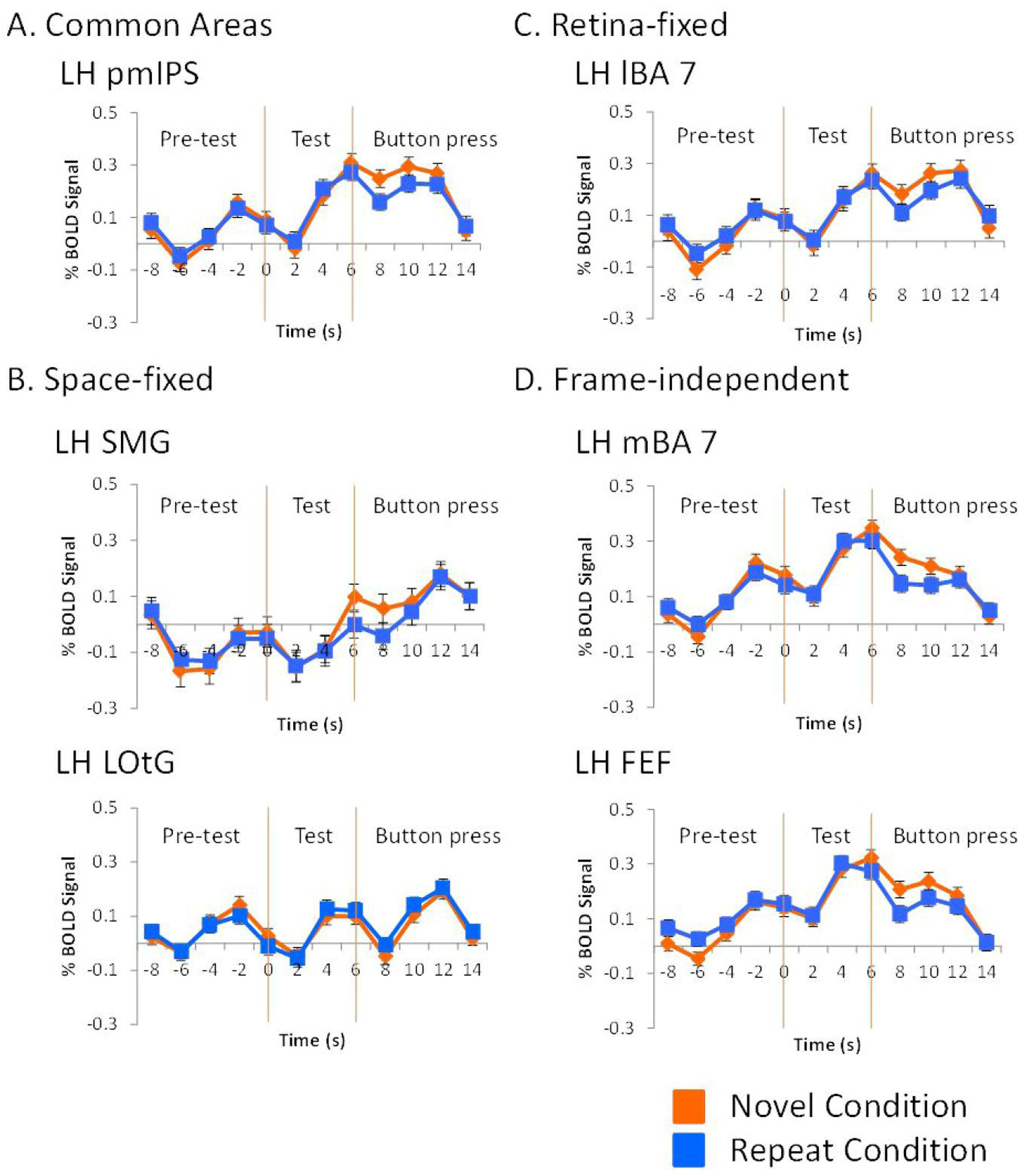
Event-related average time courses for cortical regions that show adaptation (repetition suppression or enhancement) to object orientation in areas shared by Spatially Congruent, Retinally Congruent and Spatially Incongruent conditions, as well as within each of the spatial conditions. *A*. Averaged percent BOLD signal change (%BSC) shown for the pmIPS, an area that shows adaptation among all three spatial conditions. *B*. Averaged %BSC shown for areas that show Spatially Congruent-specific adaptation, SMG and LOtG. *C*. Averaged %BSC depicted for the lBA 7, which shows Retinally Congruent-specific adaptation effects. *D*. Averaged %BSC shown for areas that demonstrate Spatially Incongruent-specific adaptation effects, mBA 7 and FEF. All event-related average time courses extracted from peak voxels within each of the areas shown. Presentation of pre-test stimulus occurs at -6 s. Test stimuli are presented at 0 s. Lastly, the button press prompt occurs at 6 s. Orange vertical lines demarcate the beginning and end of the test stimulus presentation period, and separate it from the pre-test stimulus presentation period and the button press response period. pmIPS: posterior middle Intraparietal Sulcus.

### Localizer results

Figure 8 summarizes the cortical locations of some of our main findings relative to three types of localizer. Each panel (A, B) shows activation maps plotted over an inflated brain rendering of the left hemisphere where we obtained most of our data. Only experimental areas that were near to or overlapped with the localizer data are labeled.

**Figure 8.**
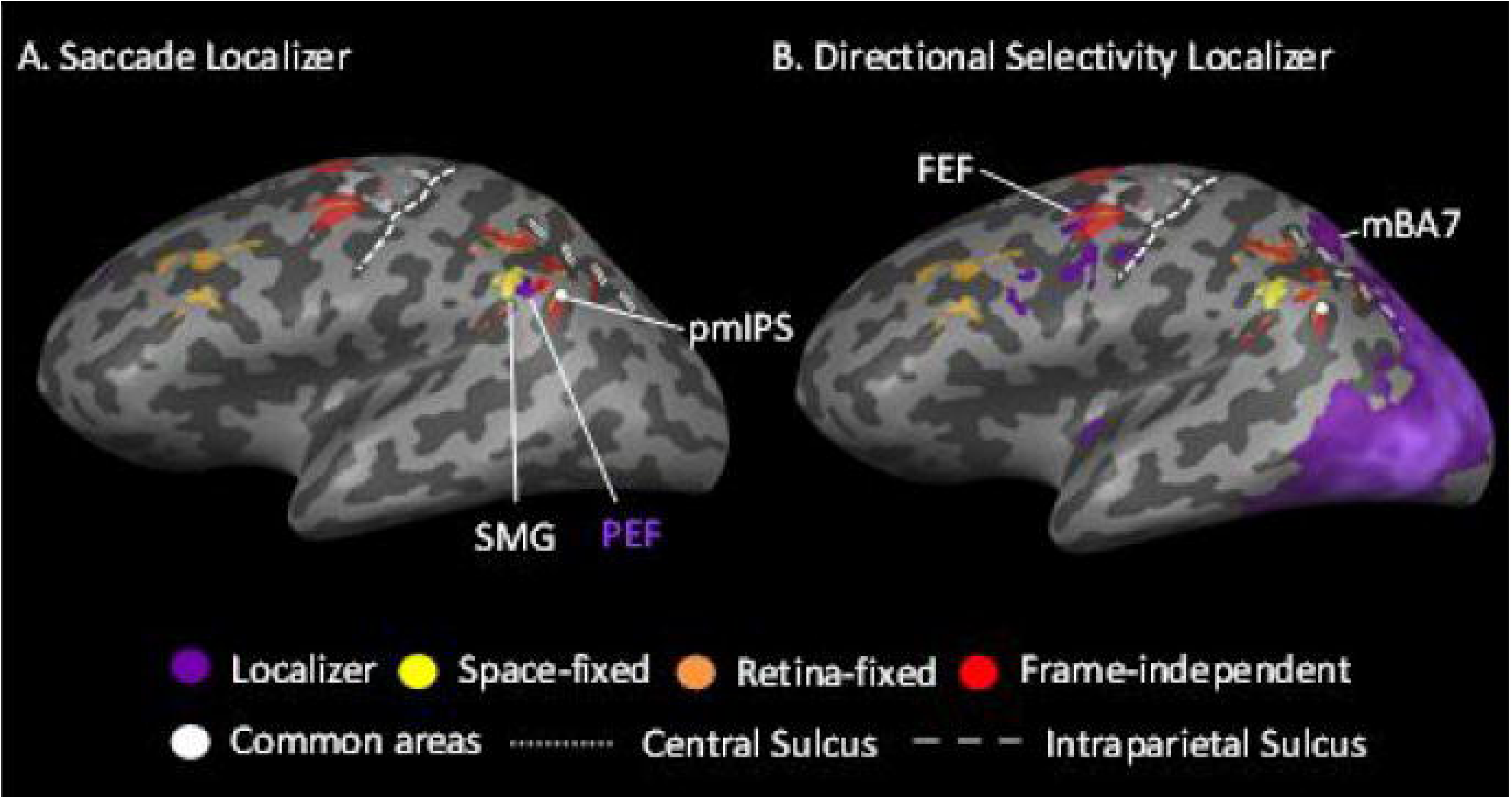
Functional localizer data for saccades and directional selectivity. ***A***. Activation map is displayed on an inflated brain of a representative participant for saccade localizer. Activation in purple depicts areas that show activation during saccade activity > fixation. ***B***. Activation map is displayed on inflated brain of representative participant for directional selectivity localizer. Using contrast for Left Visual Field > Right Visual Field in the pre-test stimulus presentation, areas that are directionally selective are denoted by the purple colour.

As shown in Figure 8A, the location of the parietal eye field (PEF) was identified by our independent saccade localizer task. Relative to the PEF, pmIPS was medially adjacent and SMG was laterally adjacent, but neither area overlapped with PEF. Finally, to determine which, if any, areas show visual field specificity within the confines of our task, we used a contrast (Adapt_LVF > Adapt_RVF) on data from the first stimulus presentation across all tasks. These data, shown in Figure 8B, were generally similar patterns to the standard retinotopy localizer, but were weighted more toward the posterior cortex. Here, mBA 7 and FEF showed partial overlap with the field-specific areas.

## Discussion

Based on the frequency of saccades in normal behavior (Rayner, 1998), the demonstrated capacity of transsaccadic memory (Irwin, 1996; Irwin & Gordon, 1998; Prime, Tsotsos, Keith & Crawford, 2007), and the ability of this system to integrate information defined in multiple frames of reference (Hayhoe et al., 1991; Melcher, 2009; Prime et al., 2006; Golomb et al., 2012), one would expect to observe extensive transfer of visual information in *the brain* during saccades. While this is generally true for location information (e.g., Duhamel et al., 1992; Medendorp et al., 2003; Merriam et al., 2003, 2007), the evidence for transsaccadic retention of object or feature information has proven to be modest or absent (Cavanagh et al., 2010; Subramanian & Colby, 2014; Lescroart et al., 2016), perhaps due to theoretical or technical difficulties in approaching this question. Here, based on the preliminary success of the fMRIa approach used by Dunkley et al. (2016), we extended this approach to again show transsaccadic feature interactions (both enhancement and suppression) in several cortical areas. Further, we found both common and different mechanisms for integrating Spatially Congruent, Retinally Congruent, and Spatially Incongruent stimuli. Overall, we found a significant difference between the patterns of activation in the Spatially Congruent condition from the other two conditions, with various detailed differences between all three patterns (to be discussed below).

Before discussing these details, it is important to note that the frame of reference (if any) used by the brain to solve a task need not be the same as the external frame of reference used to define the stimuli. For example, it is well known that the brain can use retinal coordinates to solve Spatially Congruent problems, so long as those coordinates are spatially updated relative to gaze (Duhamel et al., 1992; Medendorp et al., 2003; Merriam et al., 2003). It is also possible that object-centered information is not itself updated during saccades, but rather linked somehow to the mechanisms for spatially updating (Crawford 1997; Cavanagh et al., 2010). Thus, our three tasks (Spatially Congruent, Retinally Congruent, and Spatially Incongruent) were not designed to reveal the use of specific coordinate systems within the brain, but rather to show what networks were engaged to solve these three different transsaccadic integration tasks. Misinterpreting this fine point could potentially lead to conclusions that would contradict much of what we know about the dorsal stream visual system (Buneo & Andersen, 2006; Crawford et al., 2011; Goodale & Milner, 1992, 2008).

### Hemispheric Lateralization of Results

A striking general result of our study is that nearly all of our regions of interest were obtained in the left cortex. As discussed in more detail below, this differs from the results that Dunkley et al. (2016) obtained in their simpler ‘space-fixed’ study, but was found in all three of our spatial conditions. The general hemisphere dependencies of our results might be account/ed for by differences in the stimulus size (Busch, Debener, Kranczioch, Engel & Herrmann, 2004; Michimata & Hellige, 1987) or perhaps more likely by the greater complexity of intermingled experimental conditions in our experiment compared to the single Spatially Congruent condition in Dunkley et al. (2016). In general, the right hemisphere is associated with automatic spatial processing (Becker & Karnath, 2007; Ringman, Saver, Woolson, Clarke & Adams, 2004; Bowen, McKenna & Tallis, 1999), whereas the left hemisphere is associated with rule-based behaviors (Molfese, Freeman Jr. & Palermo, 1975; Posner, Petersen, Fox & Raichle, 1988; Harrington & Haaland, 1992; Pinker, 1998; Ullman, 2001), and possibly task switching (Corbetta & Shulman, 2002; Braver, Reynolds & Donaldson, 2003). Thus, left hemisphere may have been recruited more in the current study through some internal strategy to deal with the different stimulus combinations. Lastly, although we observed results only in the left hemisphere, we cannot draw conclusions that neurons within the right hemisphere were not at all modulated, even at sub-significant BOLD effect levels. In particular, it may be that several factors (such as rule-based and/or task-switching strategies), plus the general aspects of transsaccadic integration described below (Figure 9) had to summate in order to provide the statistically significant BOLD modulations described here.

**Figure 9.**
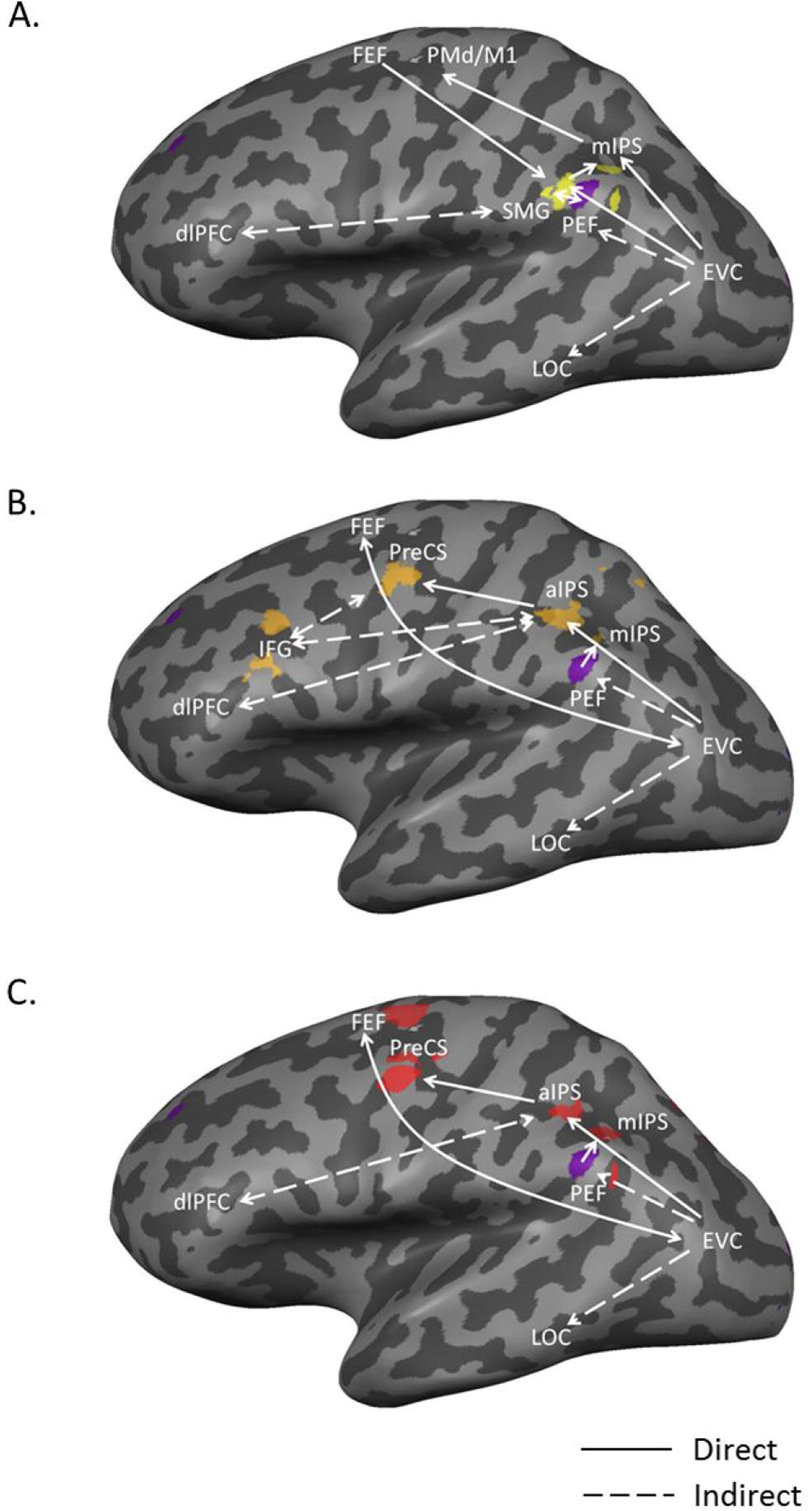
Inflated brain summary diagram of three spatial conditions and possible networks within each condition. ***A**. Spatially Congruent TSI system*. Mapped onto an inflated brain of the left hemisphere of an exemplary participant is the activation during the Spatially Congruent condition, as well as the parietal eye field (PEF; shown in purple) and possible direct and indirect networks, involving the dorsolateral prefrontal cortex (dlPFC), lateral occipitotemporal cortex (LOC, and adjacent occipital and temporal areas) and early visual cortex (EVC). ***B**. Retinally Congruent TSI system*. Shown onto an inflated brain rendering of the left hemisphere of an exemplary participant is the activation during the Retinally Congruent condition, and possible direct and indirect networks involving the PEF, EVC and dlPFC. (Motor output areas have been left out for the sake of clarity, but are assumed to also be part of this system.) ***C**. Spatially Incongruent TSI system*. Onto an inflated brain rendering of an exemplary participant is shown activation during the Spatially Incongruent condition and potential direct and indirect network connections between the activation and the PEF, EVC and dlPFC. (Motor output areas have been left out for the sake of clarity, but are assumed to also be part of this system.)

### Common areas across all three spatial conditions

In our experiment, frontal motor areas, such as pre-SMA, SMA and PMd/M1, were consistently activated in all three spatial conditions. Traditionally, these frontal motor areas have been thought of as responsible for motor processing. For example, pre-SMA and and SMA have been implicated in learning of movement sequences (Leigh & Kennard, 2005). Moreover, pre-SMA plays a role in making motor plans for effectors in response to visual information, as well as updating them based on new information (Shima, Mushiake, Saito & Tanji, 1996), whereas SMA or PMd/M1 are involved in executing manual motor plans (Grafton, Hazeltine & Ivry, 1998; Matsuzaka, Aizawa & Tanji, 1992). The coordinates for PMd/M1 are also consistent with activation in the hand motor area those found by Dechent and Frahm (2003), as well as by neuromagnetic recordings (Cheyne, Kristeva & Deecke, 1991). Thus, it seems most likely that these areas were not involved in the visual aspects of our task. Instead, they may actually reflect response-specific motor-related activity related to the button press response, which was common in all three conditions. Dunkley et al. (2016) did not report activity in these areas, but their analysis was restricted to the post-central portions of the brain.

Another common area that we observed within all three spatial conditions is an area within the intraparietal sulcus, namely pmIPS. Area pmIPS (possibly equivalent to LIP/MIP in the monkey) has been demonstrated to perform reference frame transformations related to visuomotor transformations for saccades and reach (Mullette-Gillman, Cohen & Groh, 2009, 2005; Snyder, Grieve, Brotchie & Andersen, 1998; Vesia & Crawford, 2012). Area pmIPS is just medial to the functionally localized parietal eye field (PEF), which might place it within a portion of the human ‘parietal reach region’ (Connolly, Andersen & Goodale, 2003; Gertz & Fiehler, 2015; Vesia & Crawford, 2012). This would be consistent with a role for this area in transforming information in various frames from our tasks into signals for the frontal hand motor areas described above.

However, given the proximity of the activation we found in mIPS area to our saccade localizer data, we cannot exclude the possibility that it was involved somehow in integrating saccade and feature information in all three tasks (Subramanian & Colby, 2014). This would be consistent with the role of mIPS for memory of object location in humans (Xu & Chun, 2006). This area also was not reported by Dunkley et al. (2016), but this might be accounted for by the greater complexity of our intermingled experimental tasks and the tendency for intraparietal cortex to be activated more for greater attention demands (Berhmann, Geng & Shomstein, 2004; Bisley & Goldberg, 2010; Corbetta, Kincade, Ollinger, McAvoy & Shulman, 2000; Malhotra, Coulthard & Husain, 2009).

### Novel vs. repeat modulations in the Spatially Congruent condition

Our Spatially Congruent condition was the most natural, in the sense that most naturally occuring transsaccadic stimuli are fixed in space, and thus the brain would be free to use whatever mechanisms it normally used for spatial updating and transsaccadic integration. Our Spatially Congruent condition was similar to that of Dunkley et al. (2016), with the exception that the current study used smaller stimuli (18º versus 6º), and randomly intermingled this condition with other spatial conditions. With the exception of the lateralization described above, our Spatially Congruent condition results were generally consistent with those of Dunkley et al. (2016). Both studies found repetition suppression in SMG. SMG has previously been implicated in reading (Stoeckel, Gough, Watkins & Devlin, 2009), perception of spatial orientation (Kheradmand, Lasker & Zee, 2003) and visual search (Taylor, Muggleton, Kalla, Walsh & Eimer, 2011). Inferior parietal cortex has also been associated with integrating information from the dorsal and ventral stream (Boussaoud et al., 1990; Rizzolatti & Matelli, 2003). All of these functions are consistent with the notion of transsaccadic integration of spatial and feature information. Likewise, both studies (ours and that of Dunkley et al. (2016)) found repetition enhancement in ventral occipital cortex-this region of cortex has been found to play a well-known role as part of the ventral visual stream in object recognition and identity (Kanwisher, Woods, Iacobini & Mazziotta, 1997; Malach et al., 1995; Ishai, Ungerleider, Martin, Schouten & Haxby, 1999; James, Culham, Humphrey, Milner & Goodale, 2003; Grill-Spector, Kourtzi & Kanwisher, 2001). In our task, the brain might interpret the change in stimulus to be a change in orientation of the same object, or a different object with the oppose orientation. This ambiguity may therefore be a factor that led to modulation in both SMG and LOC. Dunkley et al. (2016) accounted for the different responses in these regions (parietal RS vs. occipital RE) as the tendency of parietal cortex to respond better to Novel stimuli (Ardekani, Choi, Hossein-Zdeh, Porjesz, Tanabe, Lim et al., 2002; Linden, Prvulovic, Formisano, Voellinger, Zanella, Goebel, et al., 1999; Singh-Curry & Husain, 2009), versus occipital summation of pre- and post-saccadic feature information (Malik et al., 2015).

However, our results were not identical to those of Dunkley et al. (2016). As mentioned above, Dunkley et al. (2016) only obtained significant results in right cortex, whereas most of our significant clusters were in left cortex. In particular, Dunkley et al. (2016) found a significant transsaccadic RS effect in right SMG (with a similar trend in left SMG), whereas we only obtained a significant effect in left SMG. In addition to the general task differences described above between our studies, this might relate also to local factors. A previous study showed that TMS over right posterior parietal cortex (near both SMG and mIPS) had a greater effect than left parietal TMS on transsaccadic memory of multiple object orientations in Spatially Congruent coordinates. Right SMG has been implicated in saccades and attention to contralateral space (Perry & Zeki, 2000), whereas the left SMG has been implicated in short-term memory (Russ, Mack, Grama, Lanfermann & Knopf, 2003). All of these functions seem to relate to transsaccadic integration, so this does not appear to explain this difference in our results.

The details of our ventral stream findings also differed. Dunkley et al. (2016) reported a right extrastriate area overlapping with putative V4, whereas we localized our effect to a more lateral area (LOtG) in left cortex. In addition to falling within opposite hemispheres, these areas are thought to have different functions. V4 has recently been associated with transsaccadic updating (Neupane et al., 2016), perhaps due to re-entrant feedback from FEF (Moore & Amrstrong, 2003; Hamker, 2005; Prime et al., 2010). In contrast, LOtG is thought to be involved in recognizing gestures (Decety et al., 1997; Peigneux et al., 2000), although it is found within the lateral occipital complex (LOC) which is involved in judging the familiarity of shapes and features (Kanwisher, Woods, Iacobini & Mazziotta, 1997; Malach et al., 1995). Again, the difference in both area and laterality might have something to do with the increased spatial task complexity in our experiment and internal strategies developed by the brain to deal with this.

Finally, Dunkley et al. (2016) showed that their transsaccadic areas were not modulated by same/different features during fixation, and did not overlap with other areas that did show such modulations during fixation. We did not repeat the fixation condition here because of the need for time to do our other tasks, so we cannot assume this to be the case in our data.

### Novel vs. repeat modulations in the Retinally Congruent condition

Our study also included a Retinally Congruent fMRIa paradigm that has not previously been tested. This condition was not natural, in the sense that visual stimuli do not ordinarily move with the eye. However, it has been shown that retinal memory can be equal or superior to spatial memory (Golomb et al., 2012), perhaps because of the plethora of retina-fixed mechanisms within the visual system, some of them related to memory (Fecteau & Munoz, 2005; Golomb et al., 2008, 2010, 2011; Pratte & Tong, 2014; Talsma et al., 2013).

The unique (non-motor) areas that showed RS in the Retinally Congruent condition were lateral BA 7 and inferior frontal gyrus. BA 7 is thought to be involved in processing spatial information (Binkofski et al., 1999; Haxby et al., 1991), and our lateral area was located amongst regions running medial to the intraparietal sulcus involved in sensorimotor transformations for reach (see Vesia & Crawford 2012 for a graphic review). Area aIPS is best known for its role in grasping (Gallivan & Culham, 2015; Monaco et al., 2014; Murata, Gallese, Luppino, Kaseda & Sakata, 2000). Precentral sulcus is also associated with the skeletomotor system, although some areas show saccadic signals (Rosano et al., 2002), and our particular area overlapped with the retinotopic localizer data. Lastly, activity in the inferior frontal gyrus has been related to executive and inhibition control, which has been hypothesized to work in conjunction with the posterior parietal cortex and preSMA to respond in task-difficult and -relevant manner (Duann, Ide, Luo & Li, 2009; Hampshire, Chamberlain, Monti, Duncan & Owen, 2010).

The preceding results seem to implicate the grasp system in solving this task. This might seem odd, but the brain might tap into any available mechanism to solve an unusual task. This is not entirely out of place, because the grasp system is replete with stimulus orientation information in humans (Gallivan & Culham, 2015; Monaco et al., 2014) and macaques (Baumann, Fluet & Scherberger, 2009). In addition, multivoxel pattern analysis has recently shown that the human aIPS has a gaze-centered representation of visual targets for grasping (Leoné, Monaco, Henriques, Toni & Medendorp, 2015). And again, the use of this system might have been influenced by the behavioral output in our task: a button push. Combining these mechanisms with retinotopic mechanisms (likely most of the areas that showed up in our field-specificity localizer) would provide massive support for solving a retinotopic task.

### Novel vs. repeat modulations in the Spatially Incongruent condition

The second Novel task in this study was the Spatially Incongruent condition, perhaps the least natural condition (unless stimuli suddenly jump randomly during saccades). No condition is truly Spatially Incongruent, but in order to solve this task subjects would only have object-centered information. As described in the introduction, this would tend to predict activation of ventral stream cortical areas that utilize object-centered coordinates (Kanwisher, Woods, Iacobini & Mazziotta, 1997; Malach et al., 1995; Ishai, Ungerleider, Martin, Schouten & Haxby, 1999; James, Culham, Humphrey, Milner & Goodale, 2003; Grill-Spector, Kourtzi & Kanwisher, 2001). Alternatively, if the brain automatically tries to compare the two stimuli within a common spatial frame, whether retinal or spatial coordinates, this could actually lead to *increased* activation of dorsal stream brain areas associated with spatial location (Milner & Goodale, 1992; Ungerleider & Haxby, 1994; Culham & Kanwisher, 2001; Goodale & Westwood, 2004; Munoz, 2002; Tong, 2003). In the event, we observed a combination of these two patterns in our results, i.e., both ventral and dorsal modulations.

Consistent with the latter possibility, our Spatially Incongruent task evoked the widest area of RS effects in parietofrontal cortex, including unique areas FEF, mBA 7, as well as aIPS and mIPS and frontal precentral sulcus. We also observed RE effects in LOtG, except this time in the right hemisphere. Most of these areas we have already discussed above. FEF is famously involved in gaze control and working memory (Dias & Segraves, 1999; Goldberg & Bruce, 1990; O’Sullivan, Jenkins, Henderson, Kennard & Brooks, 1995), and is known to primarily utilize gaze-centred signals (Goldberg & Bruce, 1990; Sajad et al., 2015). FEF is thought to provide an efference copy signal of saccade motion to update space-fixed object locations (Prime et al., 2010; Duhamel et al., 1992), so this might have been suppressed by inhibitory inputs here. Conversely, subjects might have updated the first stimulus with eye position (Colby & Goldberg, 1992; Medendorp, Goltz, Vilis & Crawford, 2003; Merriam, Genovese & Colby, 2003; Nakamura & Colby, 2002) and then, transformed both stimuli into some other common coordinate frame to perform the task. Finally, it is possible that subjects used the saccade fixation points as allocentric cues to help solve this sequential task (Byrne & Crawford, 2011). Consistent with this, medial posterior parietal cortex has been associated with the use of allocentric coordinates (Uchimara, Nakano, Morito, Ando & Kitazawa, 2015). Thus, a rather diverse network of excitatory and inhibitory signals may have been required to deal with the complex and arbitrary spatial transformations involved in this task.

### Integration of our 3 networks within the general transsaccadic network

Figure 9A, B, C provides a schematic of the three *Spatially Congruent, Retinally congruent*, and *Spatially Incongruent* networks that we observed in this study and their hypothetical relationships to other brain areas that have been implicated in transsaccadic memory and integration. More specifically, the figure is summary of the cortical regions observed to be implicated within each spatial condition. Along with our results, we have also included areas that have been found to be involved in transsaccadic integration from previous studies (Prime et al., 2008; Prime et al., 2009; Tanaka et al., 2014; Malik et al., 2015). Lastly, we included links that represent possible underlying direct and indirect connections between the cortical regions we found here and the regions found in previous transsaccadic integration studies (Mishkin & Ungerleider, 1982; Sakata, Taira, Kusunoki, Murata & Tanaka, 1997; Culham & Kanwisher, 2001; James et al., 2002; Malach, Levy & Hasson, 2002; Grill-Spector et al., 2003; Goodale & Westwood, 2004). Dorsolateral prefrontal cortex has been implicated in task switching between fixation and transsaccadic memory (Tanaka et al., 2014), so we speculate that it might play a similar role here, helping to select between these networks. Early visual cortex not only provides input to the visual system in general, it has been shown to be modulated by saccade signals (McFarland, Bondy, Saunders, Cumming & Butts, 2015; Sylvester, Haynes & Rees, 2005; Ross et al., 2001) and play a role in transsaccadic spatial updating of features (Malik et al., 2015). The parietal eye fields (located amongst our 3 networks) and frontal eye fields (modulated in our Spatially Incongruent task) may provide the saccade efference copy inputs for spatial updating, so again these signals might have been suppressed or gated differently in our non-Spatially Congruent tasks. The implication is that the solution to seemingly small changes in the spatial nature of a visual task may require rather sweeping cortical network changes.

### Caveats and Limitations

One particular caveat for interpreting these data comes from the difficulty in interpreting fMRIa data. Adaptation studies encourage participants to pay attention using a task that may be related to the overall process that is being investigated (non-orthogonal task) or may be independent of what is being investigated (orthogonal task) (Grill-Spector, Kushnir, Edelman, Avidan, Itzchak et al., 1999; Gauthier, Skudlarski, Gore & Anderson, 2000; Kourtzi & Kanwisher, 2000; Winston, Henson, Fine-Goulden & Dolan, 2004). Therefore, it may be possible that cortical regions involved in attention, etc., may be found in results when analyzing data from an fMRIa study. However, subtracting the repeated condition data from the novel condition data should eliminate any activation that is found in both conditions, such as data that shows a response to the orthogonal or non-orthogonal task. It could still be worthwhile to investigate whether the results here are affected by the task utilized to engage participants.

A specific potential limitation of this study is that, in order to create these different spatial conditions, we had to present the stimulus across and within visual fields (Spatially Congruent and Incongruent conditions, and Retinally Congruent condition, respectively), as well as different eccentricities. When designing this experiment, it proved difficult to remove all of these potential confounds between our conditions without introducing other confounds (such as using larger eccentricities within one visual field). However, we made best efforts to minimize differences between tasks, given the limitations intrinsic to these spatial conditions.

Further, it is possible that participants might have experienced apparent motion when viewing the two consecutive stimuli in our tasks. However, this seems unlikely because our stimuli do not appear to meet the previously reported spatiotemporal requirements necessary to elicit apparent motion (Evans & Nettlebeck, 1993; Williams, Elfar, Eskandar, Toth & Assad, 2003; Matsuyoshi, Hirose, Mima, Fukuyama & Osaka, 2007; Larsen & Bundesen, 2009).

Lastly, the differences in overall cortical recruitment observed in our tasks (least in Spatially Congruent and most in Spatially Incongruent) might simply be explained by task difficulty. However, our behavioural results suggest that there was no difference in performance between our three spatial conditions. Further, this does not explain why some areas (such as SMG) were ‘derecruited’ in the other tasks. Therefore, it does not seem that there was a strict correlation between task difficulty and the patterns of modulation that we observed. An alternative explanation is that the less ecological the task, the more attention and processing is required to accomplish the task. In other words, the Spatially Congruent task was the most ecologically natural, whereas the Incongruent task was probably least ecologically valid.

### Conclusion: influence of spatial congruity of stimuli in transsaccadic integration

Relatively little is known about the neural mechanisms for transsaccadic integration of visual features (Melcher & Colby, 2008; Cavanagh, Hunt, Afraz & Rolfs, 2010; Prime et al., 2011). In comparison, transsaccadic updating of visual space has a well-described neurophysiological basis (Bays & Husain, 2007; Klier & Angelaki, 2008) and has been applied to understand deficits in several clinical populations (Graves & Jones, 1992; Pisella & Mattingley, 2004; Khan et al., 2005; Ansuini, Pierno, Lusher & Castiello, 2006; Ritchie, Hunt & Sahraie, 2012). We set out to test if different cortical networks are employed for transsaccadic comparisons of stimuli paired in Spatially Congruent, Retinally Congruent, and Spatially Incongruent coordinates. Clearly, this was the case. This is consistent with a recent finding that the cortical mechanisms for visually-guided reach depend on the reference frame employed in the task (Chen et al., 2014). It is noteworthy therefore that SMG, which was the most prominent unique area for the Spatially Congruent task, was no longer modulated in the other tasks, re-enforcing a special purpose for this area in real life. A general trend that we observed was the increased recruitment of parietofrontal areas as we went from the most natural (Spatially Congruent) to the most arbitrary (Spatially Incongruent) task. Since we did not observe any significant difference in performance between these three tasks, this did not appear to be due to task difficulty, but rather due to the different cognitive strategies discussed above.

## REFERENCES

1. Ardekani, B.A., Choi, S.J., Hossein-Zadeh, G.A., Porjesz, B., Tanabe, J.L., Lim, K.O., et al. (2002). Functional magnetic resonance imaging of brain activity in the visual oddball task. Cancer Biother Radio, 14, 347–356.

2. Ansuini, C., Pierno, A.C., Lusher, D., & Castiello, U. (2006). Virtual reality applications for the remapping of space in neglect patients. Restor Neurol Neurosci, 24, 431–441.

3. Averbach, E., & Coriell, A.S. (1961). Short-term memory in vision. Bell Sys Tech J, 40, 309–328.

4. Baumann, M.A., Fluet, M.-C., & Scherberger, H. (2009). Context-specific grasp movement representation in the macaque anterior intraparietal area. J Neurosci, 29, 6346–6448.

5. Bays, P.M. & Husain, M. (2007). Spatial remapping of the visual world across saccades. Neuroreport, 18, 1207–1213.

6. Becker, E., & Karnath, H.O. (2007). Incidence of visual extinction after left versus right hemisphere stroke. Stroke, 38, 3172–3174.

7. Behrmann, M., Geng, J.J., & Shomstein, S. (2004). Parietal cortex and attention. Curr Opin Neurobiol, 14, 212–217.

8. Binofski, F., Buccino, G., Posse, S., Seitz, R.J., Rizzolatti, G., & Freund, H.-J. (1999). A fronto-parietal circuit for object manipulation in man: evidence from an fMRI-study. Eur J Neurosci, 11, 3276–3286.

9. Bisley, J.W., & Goldberg, M.E. (2010). Attention, intention, and priority in the parietal lobe. Annu Rev Neurosci, 33, 1–21.

10. Bowen, A., McKenna, K., & Tallis, R.C. (1999). Reasons for variability in the reporter rate of occurrence of unilateral spatial neglect after stroke. Stroke, 30, 1196–1202.

11. Brainard, D. (1997). The psychophysics toolbox. Spat Vis, 10, 443–446.

12. Braver, T.S., Reynolds, J.R., & Donaldson, D.I. (2003). Neural mechanisms of transient and sustained cognitive control in task switching. Neuron, 39, 713–726.

13. Buneo, C.A., & Andersen, R.A. (2006). The posterior parietal cortex: sensorimotor interface for the planning and online control of visually guided movements. Neuropsychologia (Oxford), 44, 2594–2606.

14. Cavanagh, P., Hunt, A.R., Afraz, A., & Rolfs, M. (2010). Visual stability based on remapping of attention pointers. Trends Cogn Sci, 14, 147–153.

15. Chen, Y., Monaco, S., Byrne, P., Yan, X., Henriques, D.Y.P., & Crawford, J.D. (2014).Allocentric versus egocentric representation of remembered reach targets in human cortex. J Neurosci, 34, 12515-12526.

16. Cheyne, D., Kristeva, R., & Deecke, L. (1991). Homuncular organization of human motor cortex as indicated by neuromagnetic recordings. Neurosci Lett, 122, 17–20.

17. Connolly, J.D., Andersen, R.A., & Goodale, M.A. (2003). FMRI evidence for a ‘parietal reach region’ in the human brain. Exp Brain Res, 153, 140–145.

18. Corbetta, M., Kincade, J.M., Ollinger, J.M., McAvoy, M.P., & Shulman, G.L. (2000). Voluntary orienting is dissociated from target detection in human posterior parietal cortex. Nat Neurosci, 3, 292–297.

19. Corbetta, M. & Shulman, G.L. (2002). Control of goal-directed and stimulus-driven attention in the brain. Nat Neurosci Rev, 3, 201–215.

20. Crawford, J.D. 1997. Visuomotor codes for three-dimensional saccades. In: Computational and Psychophysical Mechanisms of Visual Coding, L. Harris & M. Jenkins (Eds.). Cambridge University Press, 4-102.

21. Culham, J.C., & Kanwisher, N.G. (2001). Neuroimaging of cognitive functions in human parietal cortex. Curr Opin Neurobiol, 11, 157–163.

22. Decety, J., Grezes, J., Costes, N., Perani, D., Jeannerod, M., Procyk, E., Grassi, F., & Fazio, F. (1997). Brain activity during observation of actions-influence of action content and subject’s strategy. Brain, 120, 1763–1777.

23. Dechent, P., & Frahm, J. (2003). Functional somatotopy of finger representations in human primary motor cortex. Hum Brain Mapp, 18, 272-283.

24. Dias, E.C., & Segraves, M.A. (1999). Muscimol-induced inactivation of monkey frontal eye field: effects on visually and memory-guided saccades. J Neurophysiol, 81, 2191–2214.

25. Dove, A., Pollmann, S., Schubert, T., Wiggins, C.J., & von Cramon, D.Y. (2000). Prefrontal cortex activation in task switching: an event-related fMRI study. Cogn Brain Res, 9, 103–109.

26. Duann, J.-R., Ide, J.S., Luo, X., & Li, C.-S. R. (2009). Functional connectivity delineates distinct roles of the inferior frontal cortex and presupplementary motor area in stop signal inhibition. J Neurosci, 29, 10171–10179.

27. Duhamel, J., Colby, C., & Goldberg, M. (1992). The updating of the representation of visual space in parietal cortex by intended eye movements. Science, 255, 90–92.

28. Dunkley, B.T., Baltaretu, B.R., & Crawford, J.D. (2016). Transsaccadic interactions in human parietal and occipital cortex during the retention and comparison of object orientation. Cortex, 82, 263–276.

29. Evans, G. & Nettlebeck, T. (1993). Inspection time: a flash mask to reduce apparent movement effects. Person individ Diff, 15, 91–94.

30. Fecteau, J.H., & Munoz, D.P. (2005). Correlates of capture of attention and inhibition of return across stages of visual processing. J Cog Neurosci, 17, 1714–1727.

31. Fernandez-Ruiz, J., Goltz, H.C., DeSouza, J.F.X., Vilis, T., & Crawford, J.D. (2007). Human parietal “reach region” primarily encodes intrinsic visual direction, not extrinsic movement direction, in a visual-motor dissociation task. Cereb Cortex, 17, 2283–2292.

32. Forman, S.D., Cohen, J.D., Fitzgerald, M., Eddy, W.F., Mintun, M.A., & Noll, D.C. (1995). Improved assessment of significant activation in functional magnetic resonance imaging (fMRI): use of a cluster-size threshold. Magn Reson Med, 33, 636–647.

33. Gallivan, J.P., & Culham, J.C. (2015). Neural coding within human brain areas involved in actions. Curr Opin Neurobiol, 33, 141–149.

34. Gauthier, I., Skudlarski, P., Gore, J.C., &Anderson, A.W. (2000.) Expertise for cars and birds recruits brain areas involved in face recognition. Nat Neurosci, 3, 191–197.

35. Gertz, H., & Fiehler, K. (2015). Human posterior parietal cortex encodes the movement goal in a pro-/anti-reach task. J Neurophysiol, 114, 170–183.

36. Golomb, J.D., Chun, M.M., & Mazer, J.A. (2008). The native coordinate system of spatial attention is retinotopic. J Neurosci, 28, 10654–10662.

37. Golomb, J.D., Pulido, V.Z., Albrecht, A.R., Chun, M.M., & Mazer, J.A. (2010). Robustness of the retinotopic attentional trace after eye movements. J Vis, 10, 1–12.

38. Golomb, J.D., & Kanwisher, N. (2011). Higher level visual cortex represents retinotopic, not spatiotopic, object location. Cereb Cortex doi:10.1093/cercor/bhr357.

39. Golomb, J.D., & Kanwisher, N. (2012). Retinotopic memory is more precise than spatiotopic memory. Proc Natl Acad Sci, 109, 1796–1801.

40. Goodale, M.A., & Milner, A.D. (1992). Separate visual pathways for perception and action. Trends Neurosci, 15, 20–25.

41. Goodale, M.A., & Westwood, D.A. (2004). An evolving view of duplex vision: separate but interacting cortical pathways for perception and action. Curr Opin Neurobiol, 13, 203–211.

42. Grafton, S.T., Hazeltine, E., & Ivry, R.B. (1998). Abstract and effector-specific representations of motor sequences identified with PET. J Neurosci, 18, 9420–9428.

43. Graves, R.E. & Jones, B.S. (1992). Conscious visual perceptual awareness vs. non-conscious visual spatial localisation examined with normal subjects using possible analogues of blindsight and neglect. Cogn Neuropsychol, 9, 487–508.

44. Grill-Spector, K., Henson, R., & Martin, A. (2006). Repetition and the brain: neural models of stimulus-specific effects. Trends Cogn Sci, 10, 14–23.

45. Grill-Spector, K., Kushnir, T., Edelman, S., Avidan, G., Itzchak, Y., & Malach, R. (1999). Differential processing of objects under various viewing conditions in the human lateral occipital complex. Neuron, 24, 187-203.

46. Grill-Spector, K., Kushnir, T., Hendler, T., Malach, R. (2003). The dynamics of object-selective activation coreralte with recognition performance in humans. Nat Neurosci, 3, 837–843.

47. Grill-Spector, K., & Malach, R. (2001). fMR-adaptation: a tool for studying the functional properties of human cortical neurons. Acta Psychol (Amst), 107, 293–321.

48. Hamker, F.H. (2005). The re-entry hypothesis: The putative interaction of the frontal eye field, ventrolateral prefrontal cortex, and areas V34, IT, for attention and eye movement. Cereb Cortex, 15, 431–447.

49. Hamker, F.H., Zirnksak, M., Ziesche, A., & Lappe, M. (2011). Computation models of spatial updating in peri-saccadic perception. Philos Trans R Soc Lond B Sci, 366, 554–571.

50. Hampshire, A., Chamberlain, S.R., Monti, M.M., Duncan, J., & Owen, A.M. (2010). The role of the right inferior frontal gyrus: inhibition and attentional control. Neuroimage, 50, 1313–1319.

51. Harrington, D.L., & Haaland, K.Y. (1992). Motor sequencing with left hemisphere damage: Are some cognitive deficits specific to limb apraxia? Brain, 115, 857–874.

52. Harrison, S.A., & Tong, F. (2009). Decoding reveals the contents of visual working memory in early visual areas. Nature, 458, 632–635.

53. Hayhoe, M., Lachter, J., & Feldman, J. (1991). Integration of form across saccadic eye movements. Perception (London), 20, 393–402.

54. Haxby, J.V., Grady, C.L., Horwitz, B., Ungerleider, L.G., Mishkin, M., Carson, R.E., Herscovitch, P., Schapiro, M.B., & Rapoport, S.I. (1991). Dissociation of spatial and object visual processing pathways in human extrastriate cortex. Proc Nad Acad Sci USA, 88, 1621–1625.

55. Irwin, D.E. (1992). Memory for position and identity across eye movements. J Exp Psychol Learn Mem Cogn, 18, 307-317.

56. Irwin, D.E. (1996). Integrating information across saccadic eye movements. Curr Dir Psychol Sci, 5, 94–100.

57. Irwin, D.E., & Gordon, R.D. (1998). Eye movements, attention and transsaccadic memory. Vis Cog, 5, 127–155.

58. Ishai, A., Ungerleider, L.G., Martin, A., Schouten, J.L., & Haxby, J.V. (1999). Distributed representation of object in the human ventral visual pathway. Proc Nad Acad Sci USA, 96, 9379–9384.

59. James T.W., Humphrey, G.K., Gati, J.S., Menon, R.S., Goodale, M.A (2002). Differential effects of viewpoint on object-driven activation in dorsal and ventral streams. Neuron, 35, 79-801.

60. James, T.W., Culham, J., Humphrey, G.K., Milner, A.D., & Goodale, M.A. (2003). Ventral occipital lesions impair object recognition but not object-directed grasping; an fMRI study. Brain, 126, 2463–2475.

61. James, T.W., & Gauthier, I. (2006). Repetition-induced changes in BOLD response reflect accumulation of neural activity. Hum Brain Mapp, 27, 37–46.

62. Johnson, J.S., Hollingworth, A., & Luck, S.J. (2008). The role of attention in the maintenance of feature bindings in visual short-term memory. J Exp Psychol Hum Percept Perform, 34, 41–55.

63. Kanwisher, N., Woods, R.P., Iacoboni, M., & Mazziotta, J.C. (1997). A locus in human extrastriate cortex for visual shape analysis. J Cognit Neurosci, 9, 133–142.

64. Khan, A.Z., Pisella, L., Vighetto, A., Cotton, F., Luauté, J., Boisson, D., et al. (2005). Optic ataxia errors depend on remapped, not viewed, target location. Nat Neurosci, 8, 418–420.

65. Kheradmand, A., Lasker, A., & Zee, D.S. (2013). Transcranial magnetic stimulation (TMS) of the supramarginal gyrus: a window to perception of upright. Cereb Cortexdoi:10.1093/cercor/bht267.

66. Klier, E., & Angelaki, D. (2008). Spatial updating and the maintenance of visual constancy. Neuroscience, 156, 801–818.

67. Kourtzi, Z. & Kanwisher, N. (2000). Cortical regions involved in perceiving object shape. J Neurosci, 20, 3310–3318.

68. Krekelberg, B., Boynton, G.M., & van Wezel, R.J.A. (2006). Adaptation: from single cells to BOLD signals. Trends Neurosci, 29, 250–256.

69. Kriegeskorte, N., Lindquist, M.A., Nichols, T.E., Poldrack, R.A., & Vul, E. 2010. Everything you ever wanted to know about cirucular analysis, but were afraid to ask. J Cereb Blood Flow Metab. doi:10.1038/jcbfm.2010.86

70. Larsen, A., & Bundesen, C. (2009). Common mechanisms in apparent motion perception and visual pattern matching. Scand J Psychol, 50, 526–534.

71. Leigh, R.J., & Kennard, C. (2004). Using saccades as a research tool in the clinical neurosciences. Brain, 127, 460–477.

72. Leoné, F., Monaco, S., Henriques, Y.P.D., Toni, I., & Medendorp, P.W. (2015). Flexible reference frames for grasp planning in human parieto-frontal cortex. eNeuro. doi:10.1523/eneuro.0008-15.2015.

73. Linden, D.E.J., Prvulovic, D., Formisano, E., Voellinger, M., Zanella, F.E., Goebel, R., et al. (1999). The functional neuroanatomy of target detection: An fMRI study of visual and auditory oddball tasks. Cereb Cortex, 9, 815–823.

74. Logothetis, N.K., Paul,s J., & Poggio, T. (1995). Shape representation in the inferior temporal cortex of monkeys. Curr Biol, 5, 552–563.

75. Malach, R., Levy, I., Hasson, U. (2002). The topography of high-order human object areas. Trends Cog Sci, 6,176–184.

76. Malach, R., Reppas, J.B., Kwong, K.K., Jiang, H., Kennedy, W.A., Ledden, P.J., Brady, J., Rosen, B.R., & Tootell, R.B. (1995). Object-related activity revealed by functional magnetic resonance imaging in human occipital cortex. Proc Natl Acad Sci USA, 92, 8135–8138.

77. Malhotra, P., Coutlhard, E.J., & Husain, M. (2009). Role of right posterior parietal cortex in maintaining attention to spatial locations over time. Brain. DOI:http://dx.doi.org/10.1093/brain/awn350

78. Malik, P., Dessing, J.C., & Crawford, J.D. (2015). Role of early visual cortex in transsaccadic memory of object features. J Vis, 15, 1–7.

79. Matsuyoshi, D., Hirose, N., Mima, T., Fukuyama, H., & Osaka, N. (2007). Repetitive transcranial magnetic stimulation of human MT+ reduces apparent motion perception. Neurosci Lett, 429, 131–135.

80. Matsuzaka, Y., Aizawa, H., & Tanji, J. (1992). A motor area rostral to the supplementary motor area (presupplementary motor area) in the monkey: neuronal activity during a learned motor task. J Neurophysiol, 68, 653–662.

81. McFarland, J.M., Bondy, A.G., Saunders, R.C., Cumming, B.G., & Butts, D.A. (2015). Saccadic modulation of stimulus processing in primary visual cortex. Nat Commun 6: doi:10.1038/ncomms9110.

82. Medendorp, W.P., Goltz, H.C., Vilis, T., & Crawford, J.D. (2003). Gaze-centered updating of visual space in human parietal cortex. J Neurosci, 23, 6209–6214.

83. Melcher, D. (2009). Selective attention and the active remapping of object features in transsaccadic perception. Vision Res, 49, 1249-1255.

84. Melcher, D., & Colby, C.L. (2008). Spatiotopic transfer of visual-form adaptation across saccadic eye movements. Curr Biol, 15, 1745–1748.

85. Merriam, E.P., Genovese, C.R., & Colby, C.L. (2003). Spatial updating in human parietal cortex. Neuron, 39, 361–373.

86. Merriam, E.P., Genovese, C.R., & Colby, C.L. (2007). Remapping in human visual cortex. J Neurophysiol, 97, 1738–1755.

87. Michimata, C., & Hellige, J.B. (1987). Effects of blurring and stimulus size on the lateralized processing of nonverbal stimuli. Neuropsychologia, 25, 397–407.

88. Milner, A.D., & Goodale, M.A. (2008). Two visual systems re-viewed. Neuropsychologia (Oxford), 46, 774–785.

89. Mishkin, M. & Ungerleider, L.G. (1982). Contribution of striate inputs to the visuospatial functions of parieto-preoccipital cortex in monkeys. Behav Brain Res, 6, 57–77.

90. Molfese, D.J., Freeman, Jr., R.B., & Palermo, D.S. (1975). The ontogeny of brain lateralization for speech and nonspeech stimuli. Brain Lang, 2, 356–368.

91. Monaco, S., Chen, Y., Medendorp, W.P., Crawford, J.D., Fiehler, K., & Henriques, D.Y.P. (2014). Functional magnetic resonance imaging adaptation reveals the cortical networks for processing grasp-relevant object properties. Cereb Cortex doi:10.1093/cercor/bht006.

92. Moore, T., & Armstrong, K.M. (2003). Selective gating of visual signals by microstimulation of frontal cortex. Nature, 421, 370–373.

93. Motter, B.C. (1994). Neural correlates of feature selective memory and pop-out in extrastriate area V4. J Neurosci, 14, 2190-2199.

94. Mullette-Gillman, O.A., Cohen, Y.E., & Groh, J.M. (2005). Eye-centered, head-centered, and complex coding of visual and auditory targets in the intraparietal sulcus. J Neurophysiol, 94, 2331–2352.

95. Mullette-Gillman, O.A., Cohen, Y.E., & Groh, J.M. (2009). Motor-related signals in the intraparietal cortex encode locations in a hybrid, rather than eye-centered reference frame. Cereb Cortex, 19, 1761–1775.

96. Munoz, D.P. (2002). Commentary: saccadic eye movements: overview of neural circuitry. Prog Brain Res, 40, 89–96.

97. Murata, A., Gallese, V., Luppino, G., Kaseda, M., & Sakata, H. (2000). Selectivity for the shape, size, and orientation of objects for grasping in neurons of monkey parietal area AIP. J Neurophysiol, 83, 2580–2601.

98. Nakamura, K., & Colby, C.L. (2002). Updating of the visual representation in monkey striate and extrastriate cortex during saccades. Proc Natl Acad Sci, 99, 4026–4031.

99. Neupane, S., Guitton, D., & Pack, C.C. (2016). Two distinct types of remapping in primate cortical area V4. Nat Commun. doi:10.1038/ncomms10402.z

100. O’Sullivan, E.P., Jenkins, I.H., Henderson, L., Kennard, C., & Brooks, D.J. (1995). The functional anatomy of remembered saccades: a PET study. Neuroreport (Oxford), 6, 2141–2144.

101. Peigneux, P., Salmon, E., van der Linden, M., Garraux, G., Aerts, J., Delfiore, G., Degueldre, C., Luxen, A., Orban, G., & Franck, G. (2000). The role of the lateral occipitotemporal junction and area MT/V5 in the visual analysis of upper-limb postures. NeuroImage, 11, 644–655.

102. Perry, R.J., & Zeki, S. (2000). The neurology of saccades and covert shifts in spatial attention. Brain, 123, 2273–2288.

103. Pigott, S., & Milner, B. (1993). Memory for different aspects of complex visual scenes after unilateral temporal- or frontal-lobe resection. Neuropsychologia (Oxford), 31, 1–15.

104. Pinker, S. (1997). Words and rules in the human brain. Nature, 387, 547-548.

105. Pisella, L. & Mattingley, J.B. The contribution of spatial remapping impairments to unilateral visual neglect. Neurosci Biobehav Rev, 28, 181–200.

106. Posner, M.I., Petersen, S.E., Fox, P.T., & Raichle, M.E. (1988). Localization of cognitive operations in the human brain. Science, 240, 1627–1631.

107. Poth, C., Herwig, A., & Schneider, W.X. (2015). Breaking object correspondence across saccadic eye movements deteriorates object recognition. Front Syst Neurosci, 9, doi:10.3389/fnsys.2015.00176.

108. Poth, C. & Schneider, W.X. (2016). Breaking object correspondence across saccades impairs object recognition: The role of color and luminance. J Vis, 16, doi:10.1167/16.11.1.

109. Pouget, P., Emeric, E.E., Stuphorn, V., Reis, K., & Schall, J.D. (2005). Chronometry of visual responses in frontal eye field, supplementary eye field, and anterior cingulate cortex. J Neurophysiol, 94, 2086–2092.

110. Pratte, M.S., & Tong, F. (2014). Spatial specificity of working memory representations in the early visual cortex. J Vis, 14, 1–12.

111. Prime, S.L., Niemeier, M., & Crawford, J.D. (2006). Transsaccadic integration of visual features in a line intersection task. Exp Brain Res, 169, 532–548.

112. Prime, S.L., Tsotsos, L., Keith, G.P., & Crawford, J.D. (2007). Visual memory capacity in transsaccadic integration. Exp Brain Res, 180, 609–628.

113. Prime, S.L., Vesia, M., & Crawford, J.D. (2008). Transcranial magnetic stimulation over posterior parietal cortex disrupts transsaccadic memory of multiple objects. J Neurosci, 28, 6938–6949.

114. Prime, S.L., Vesia, M., & Crawford, J.D. (2009). TMS over human frontal eye fields disrupts transsaccadic memory of multiple objects. Cereb Cortex, 20, 759–772.

115. Prime, S.L., Vesia, M., & Crawford, J.D. (2011). Cortical mechanisms for transsaccadic memory and integration of multiple object features. Phil Trans R Soc B, 366, 540–553.

116. Ringman, J.M., Saver, J.L., Woolson, R.F., Clarke, W.R., & Adams, H.P. (2004). Frequency, risk factors, anatomy, and course of unilateral neglect in an acute stroke cohort. Neurology, 63, 468–474.

117. Ritchie, K.L., Hunt, A.R., & Sahraie, A. (2012). Trans-saccadic priming in hemianopia: Sighted-field sensitivity is boosted by a blind-field prime. Neuropsychologia, 50, 997–1005.

118. Rosano, C., Krisky, C.M., Welling, J.S., Eddy, W.F., Luna, B., Thulborn, K.R., & Sweeney, J.A. (2002). Pursuit and saccadic eye movement subregions in human frontal eye field: a high-resolution fMRI investigation. Cereb Cortex, 12, 107–115.

119. Ross, J., Morrone, M.C., Goldberg, M.W., & Burr, D.C. (2001). Changes in visual perception at the time of saccades. Trends Neurosci, 24, 113–121.

120. Russ, M.O., Mack, W., Grama, C.-R., Lanfermann, H., & Knopf, M. (2003). Enactment effect in memory: evidence concerning the function of the supramarginal gyrus. Exp Brain Res, 149, 497–504.

121. Sajad, A., Sadeh, M., Keith, G.P., Yan, X., Wang, H., & Crawford, J.D. (2014), Visual-motor transformation within frontal eye fields during head-unrestrained gaze shifts in the monkey. Cereb Cortex doi:10.1093/cercor/bhu279.

122. Sakata H, Taira M, Kusunoki M, Murata A, Tanaka Y (1997) The parietal association cortex in depth perception and visual control of hand action. Trends Neurosci, 20, 350–357.

123. Shima, K., Mushiake, H., Saito, N., & Tanji, J. (1996). Role for cells in the presupplementary motor area in updating motor plans. Proc Natl Acad Sci 93, 8694–8698.

124. Snyder, L.H., Grieve, K.L., Brotchie, P., & Andersen, R.A. (1998). Separate body- and world-referenced representations of visual space in parietal cortex. Nature, 394, 887–891.

125. Stoeckel, C., Gough, P.M., Watkins, K.E., & Devlin, J.T. (2009). Supramarginal gyrus involvement in visual world recognition. Cortex, 45, 1091–1096.

126. Subramanian, J., & Colby, C.L. (2014). Shape selectivity and remapping in dorsal stream visual area LIP. J Neurophysiol, 111, 613–627.

127. Sylvester, R., Haynes, J.-D., & Rees, G. (2005). Saccades differentially modulate human LGN and V1 responses in the presence and absence of visual stimulation. Curr Biol, 15, 37–41.

128. Talairach, J., & Tournoux, P. (1988). Co-planar stereotaxic atlas of the human brain. New York: Thieme Medical.

129. Talsma, D., White, B.J., Mathot, S., Munoz, D.P., & Theeuwes, J. (2013). A retinotopic attentional trace after saccadic eye movements: evidence from event-related potentials. J Cog Neurosci, 25, 1563–1577.

130. Tanaka, L.L., Dessing, J.C., Malik, P., Prime, S.L., & Crawford, J.D. (2014). The effects of TMS over dorsolateral prefrontal cortex on transsaccadic memory of multiple objects. Neuropsychologia (Oxford), 63, 185–193.

131. Taylor, P.C.J., Muggleton, N.G., Kalla, R., Walsh, V., & Eimer, M. (2011). TMS of the right angular gyrus modulates priming of pop-out in visual search: combined TMS-ERP evidence. J Neurophysiol, 106, 3001–3009.

132. Tong, F. (2003). Primary visual cortex and visual awareness. Nat Neurosci 4, 219–229.

133. Turi, M., & Burr, D. (2012). Spatiotopic perceptual maps in humans: evidence from motion adaptation. Proc R Soc B, 279, 3091–3097.

134. Uchimara, M., Nakano, T., Morito, Y., Ando, H., & Kitazawa, S. (2015). Automatic representation of a visual stimulus relative to a background in the right precuneus. Eur J Neurosci, 42, 1651–1659.

135. Ullman, M.T. (2001). A neurocognitive perspective on language: The declarative/procedural model. Nat Rev Neurosci, 2, 717–726.

136. Ungerleider, L.G., & Haxby, J.V. (1994). ’What’ and ’where’ in the human brain. Curr Opin Neurobiol, 4, 157–165.

137. Vesia, M., & Crawford, J.D. (2012). Specialization of reach function in human posterior parietal cortex. Exp Brain Res, 221, 1–18.

138. Williams, Z.M., Elfar, J.C., Eskandar, E.N., Toth, L.J., & Assad, J.A. (2003). Parietal activity and the perceived direction of ambiguous apparent motion. Nat Neurosci, 6, 616–623.

139. Winston, J.S., Henson, R.N.A., Fine-Goulden, M.R., & Dolan, R.J. (2004). fMRI-adaptation reveals dissociable neural representations of identity and expression in face perception. J Neurophysiol, 92, 1830–1839.

140. Wolf, C., & Schütz, A.C. (2015). Transsaccadic integration of peripheral and foveal feature information is close to optimal. J Vis 16(16): 11–18

141. Xu, Y., & Chun, M.M. (2006). Dissociable neural mechanisms supporting visual short-term memory for objects. Nature, 440, 91–95.

